# Capturing *in situ* Virus-Host Range and Interaction Dynamics through Gene Fusion with epicPCR

**DOI:** 10.1101/2020.08.14.250803

**Authors:** Eric G. Sakowski, Keith Arora-Williams, Funing Tian, Ahmed A Zayed, Olivier Zablocki, Matthew B. Sullivan, Sarah P. Preheim

## Abstract

Viruses impact microbial diversity, phenotype, and gene flow through virus-host interactions that in turn alter ecology and biogeochemistry. Though metagenomics surveys are rapidly cataloging viral diversity, capturing specific virus-host interactions *in situ* would identify hosts for novel viruses and reveal influential ecological or environmental factors. We leveraged metagenomics and a high-throughput, cultivation-independent gene fusion technique (epicPCR) to investigate viral diversity and virus-host interactions over time in a critical estuarine environment, the Chesapeake Bay. EpicPCR captured *in situ* virus-host interactions for viral clades with no closely related database representatives. Abundant freshwater Actinobacteria lineages were the most common hosts for these poorly characterized viruses, and observed viral interactions with one abundant Actinobacterial population (*Rhodoluna*) were correlated with environmental factors. Tracking virus-host interaction dynamics also revealed ecological differences between multi-host (generalist) and single-host (specialist) viruses. Generalist viruses had significantly longer periods with observed virus-host interactions but specialist viruses were observed interacting with hosts at lower minimum abundances, suggesting more efficient interactions. Together, these observations reveal ecological differences between generalist and specialist viruses that provide insight into evolutionary trade-offs. Capturing *in situ* interactions with epicPCR revealed environmental and ecological factors that shape virus-host interactions, highlighting epicPCR as a scalable new tool in viral ecology.

## Introduction

Viruses impact the diversity and function of microbial communities from the open ocean^1^ and soils^2^ to the human gut^3^. Bulk measurements implicate viruses as major contributors to daily bacterial mortality (20-40% in surface waters^4^), carbon export^5^, and biogeochemical cycling^6^. In the Chesapeake Bay, the largest estuary in the United States, high viral production rates (∼8 × 10^6^ viruses ml^-1^ h^-1^) suggest up to 20% of the bacterial community can be lysed by viruses per hour,^7^ although environmental factors, such as tidal mixing of fresh and marine water, could modulate viral production and viral-mediated mortality, as observed in other ecosystems ^8^. These community-level impacts are the aggregate of a myriad of individual virus-host infections that result in an estimated 10^23^ infections per second in the oceans worldwide^9^. However, cultivation^10,11^ and theoretical models^12,13^ suggest viral pressure is not equally distributed across the microbial community, implying a subset of virus-host infections could contribute disproportionately to microbial mortality with implications for microbial community diversity and biogeochemical cycling. Since individual virus-host interactions (i.e. any physical association that could lead to an active infection including attachment, injection of genetic material, latent period, lysogeny) could be influenced by different ecological and environmental factors, establishing connections between virus-host interactions and environmental factors will be the first step in fine-tuning ecosystem models to better predict variability in viral productivity and bacterial mortality.

Viral metagenomics has greatly expanded our knowledge of environmental viral diversity; yet, linking these viruses to their hosts remains a bottleneck for investigating the relative ecosystem impact of individual virus-host pairs. Bioinformatics approaches applied to bacterial and viral shotgun metagenomes show promise for identifying hosts for viral communities. A recent study linked 35% of shotgun-assembled soil viral populations to putative hosts through a combination of bioinformatics approaches for host identification, such as CRISPR spacer matches, sequence homology, and k-mer frequencies^2^. Recent floods of oceanic virus data^14^ and metagenome-assembled genomes (MAGs)^15^ will likely improve *in silico* host predictions for marine viruses. Still, it is estimated that only 10% of uncultivated bacterial lineages contain CRISPR-Cas systems^16^, limiting efforts to link viruses and hosts by CRISPR spacer homology. Comparisons of mycobacteriophage indicate some viruses may switch hosts too quickly for their genomes to evolve similar DNA signatures to their host^17^, thus constraining *in silico* predictions via k-mer frequencies. Other techniques, like spot, plaque and liquid assays^18^ and viral tagging^19,20^ require cultivation of the host, preventing investigations of viral interactions for the majority of bacterial populations that cannot be readily cultivated. Cultivation-independent approaches like the microscopy-based phageFISH^21^ and microfluidic digital polymerase chain reaction (PCR)^22^ either require laborious and database-dependent probe design and optimization or fail to scale due to few micro-chambers yielding positive interactions, respectively. Furthermore, these methods cannot identify virus-host interaction dynamics or the ecological and environmental factors that influence these interactions. Thus, there is a need for new, scalable experimental methods that capture virus-host interactions *in situ*.

Here we sought to investigate the ecology of viruses interacting with key bacterial populations in the Chesapeake Bay. We leveraged viral metagenomics to study viral diversity and seasonal dynamics at population-scale resolution^19^. Furthermore, we repurposed emulsion paired isolation-concatenation PCR (epicPCR)^23^ to evaluate *in situ* virus-host interaction dynamics of ecologically important bacterial populations. This analysis revealed that abundant freshwater Actinobacteria populations were the primary host of poorly characterized viruses sharing genetic similarity with cyanosipho-and-podoviruses. Additionally, environmental factors partially explained the observed interactions between one abundant Actinobacterial population (*Rhodoluna*) and viral clades. We also identified ecological differences (i.e. observed “interaction life-span” and minimum host abundance) between specialist and generalist viruses. These results demonstrate that this approach can be applied as a scalable and complementary new tool in viral ecology.

## Results

### Chesapeake Bay viral populations are novel, endemic and seasonally dynamic

To investigate temporal changes in viral populations and virus-host interactions, water samples were collected from May 2017 and May to December 2018 in the Rhode River, a tidal estuary on the Western Shore of the Chesapeake Bay (Fig. S1). We employed short-read shotgun metagenomics to the bacterial fraction and short-/long-read hybrid metagenomic approaches^24^ to the viral fraction of samples to investigate bacterial and viral diversity, respectively. In total, 9,392 viral populations (approximately species level genotypes dereplicated using 95% ANI over 80% of the contig; size > 5 kb^14,25-27^) were assembled from the metagenomic libraries – 7,801 from the viral fraction and 1,591 from microbial-fraction (> 0.2 µm) metagenomes. Viral diversity in the Chesapeake Bay was distinct from reference database representatives. Chesapeake Bay viral populations formed 473 clusters in a network analysis of shared genes^28^, only 40 (8%) of which also contained RefSeq representatives. Over 2,000 additional Chesapeake Bay viral populations were not assigned a single cluster. Chesapeake Bay viral populations also shared little overlap with the Global Ocean Virome 2.0 (GOV 2.0) dataset^14^ (approximately 1% of the viral populations could be detected in any GOV2.0 sample) and other estuarine samples^29-31^ even after accounting for differences in sequencing depth between samples (Fig. 1). This highlights the novelty and potentially endemic nature of Chesapeake Bay viral populations and is consistent with prior inferences of endemism from cyanophage gene markers^32^. This endemism is likely attributable to the Bay’s long residence time (6-7 months) promoting the development of distinctly estuarine microbial populations^33^.

**Figure 1.**
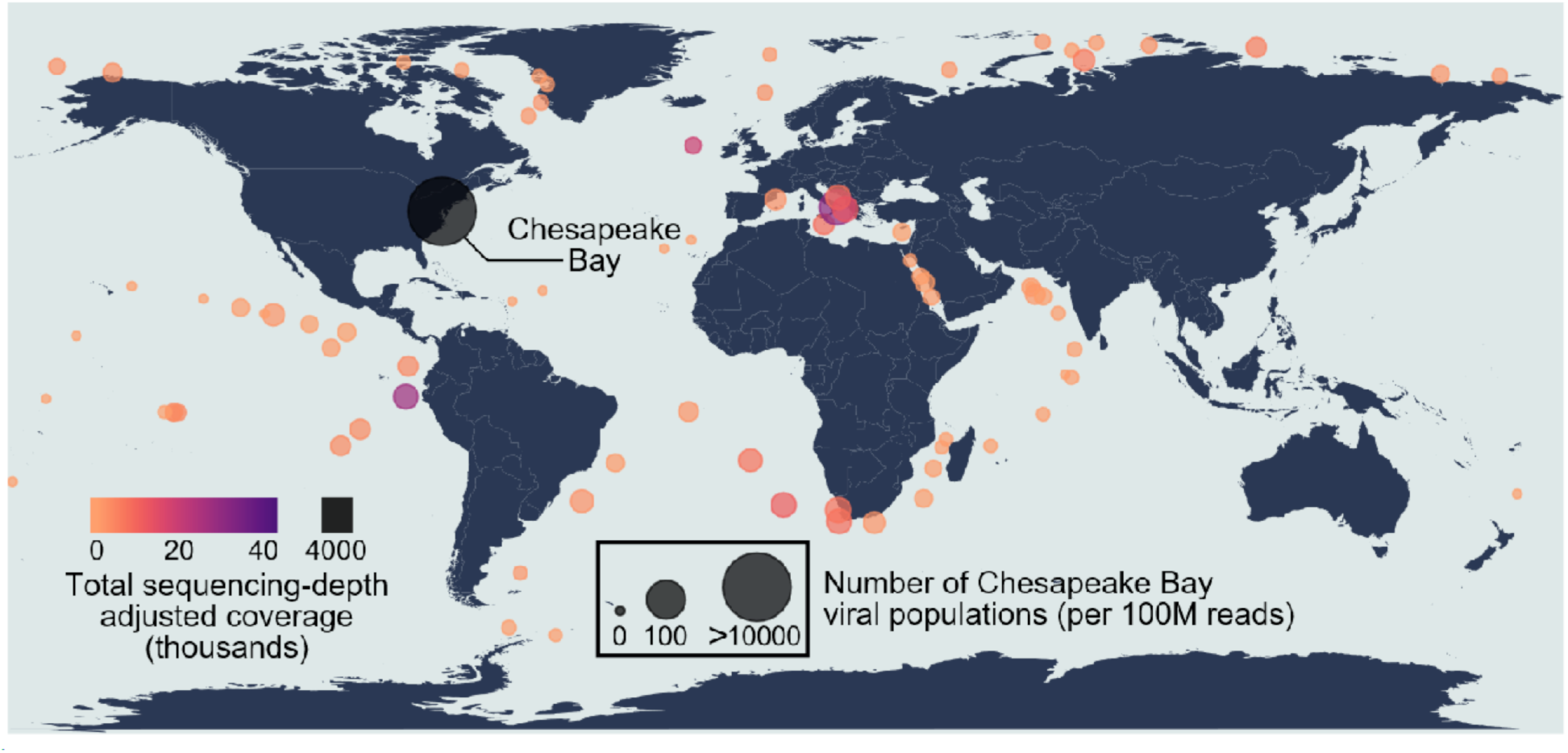
Global distribution and abundance of Chesapeake Bay viral populations. The Chesapeake Bay viral populations were highly endemic as less than 100 populations were shared with any other location in the global ocean viromes (GOV2.0) dataset (circle size) and recruited significantly less reads from the GOV2.0 stations (circle color). A viral population was considered detected in a GOV2.0 sample if the sample’s reads covered at least 70% of the genomic length of the viral population. Due to differences in the sequencing depth between libraries, the number of shared viral populations and their total coverage (tpmean; see Materials and Methods) were adjusted for each library size. For the stations that have multiple samples, the maximum value is reported.

Consistent with previous reports from the Chesapeake Bay ^34,35^, bacterial and viral communities displayed distinct seasonal trends (Fig. 2). Winter viral communities from 2018 shared nearly three times more viral populations with a winter 2012 sample^36^ than with spring and summer samples from the same year (i.e. 2018; Figs. 2B, S2). Likewise, spring 2017 and 2018 samples shared half of their viral populations with each other but fewer than 30% of their viral populations with other seasons in 2018 (Figs. 2B, S2), illustrating that Chesapeake Bay viral communities had greater seasonal than inter-annual variability. The number of shared populations between samples was similar whether comparing across entire viral contigs or a single gene. Ribonucleotide reductase (RNR) genes, which are broadly distributed across aquatic dsDNA lytic viruses^37^, were extracted from the > 5 kb viral contigs and clustered at 95% nucleotide identity. The number of shared viral populations between samples derived from this single gene was highly correlated with the number of shared populations using entire contigs (r^2^ = 0.95, p < 0.001; Fig. S3), marking RNRs as a good proxy of aquatic viral diversity in our study.

**Figure 2.**
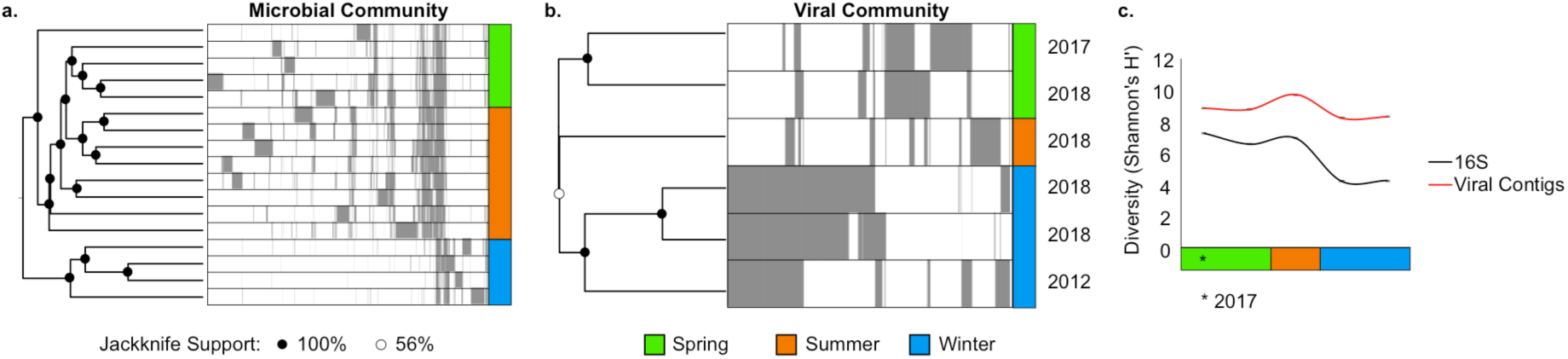
Seasonal dynamics of Chesapeake Bay bacterial and viral communities. A) Hierarchical clustering of bacterial communities based on 16S rRNA gene sequences shows that microbial communities are most similar within the same season. Libraries were sub-sampled at 150,000 observations with ten jackknifed replicates. Individual sequence variants are colored gray. B) Hierarchical clustering of viral communities from viral populations > 5kb collected across different years (see Materials and Methods) shows that seasonal community similarity exceeds within-year community similarity. Libraries were sub-sampled at 300,000 observations. Individual viral populations are colored gray. For the deeply-sequenced 2012 sample, only viral populations that were observed in the shallower 2017-2018 samples were included for clarity. C) Bacterial and viral diversity (Shannon’s *H*’ index) for paired 16S rRNA gene and viral metagenomes were significantly correlated (r^2^ = 0.6, p < 0.05). Viral metagenomes were subsampled to the smallest metagenome (15 M reads) prior to assembly for all Shannon’s *H’* diversity analyses. Diversity indices represent an average of ten jackknifed replicates at a sampling depth of 10,000 observations. Error bars are SD.

### Actinobacteria are commonly identified hosts for observed viral populations

To determine the bacterial hosts for observed viral populations, we used both bioinformatics and experimental approaches. First, we inferred hosts associated with observed viral RNR genes through similarity to reference sequences with known bacterial hosts. RNR genes observed in the spring most frequently shared homology with myoviruses that infect Cyanobacteria and Gammaproteobacteria, while winter samples showed an increase in podoviruses infecting Alphaproteobacteria. August samples were largely composed of populations with no viral reference database representatives (Fig. S4a), and overall 54 ± 19% of the RNR sequences were not classified using this homology-based approach. However, even ‘classifiable’ RNR sequences were often only distantly related to viral reference sequences (Fig. S4b), and homology to viral references does not necessarily indicate a common host. For example, Pelagibacter phage HTVC008M and T4-like cyanophages share 71% RNR nucleotide identity and would be classified together by our cutoffs; yet, they infect hosts from different phyla. Therefore, gene homology may aid predictions of phage morphology but lacks the resolution to predict hosts for environmental viruses that are only distantly related to reference database representatives.

To further identify hosts associated with viral RNR sequences, we modified an existing single-cell gene fusion protocol (epicPCR)^23^ and applied it to a large set of samples from our study site (Fig. S1b). EpicPCR takes advantage of the close physical proximity of phage and host DNA within infected bacterial cells to fuse viral marker and host 16S ribosomal RNA (rRNA) genes within emulsion droplets, providing single-cell level resolution (Fig. 3). The method requires no *a priori* knowledge of putative hosts. However, because viruses lack a universal marker gene^38^, viral diversity assessed by epicPCR is limited by viral marker gene choice. Based on reference genomes, cyanosipho-and-podoviral RNR genes form a distinct, monophyletic clade (the Cyano SP clade^39^), which enabled the development of primers to target these viral populations by epicPCR. Furthermore, cyanopodoviruses were shown to be the dominant cyanophage populations in the Chesapeake Bay in previous studies^34^; yet, cyanosipho-and-podoviral RNR genes from the Chesapeake Bay viral metagenomes were only distantly related to reference cyanophage sequences (Fig. S4b). Therefore, Cyano SP RNR was an ideal marker gene choice for investigating poorly characterized cyanophage-like diversity in this system.

**Fig. 3.**
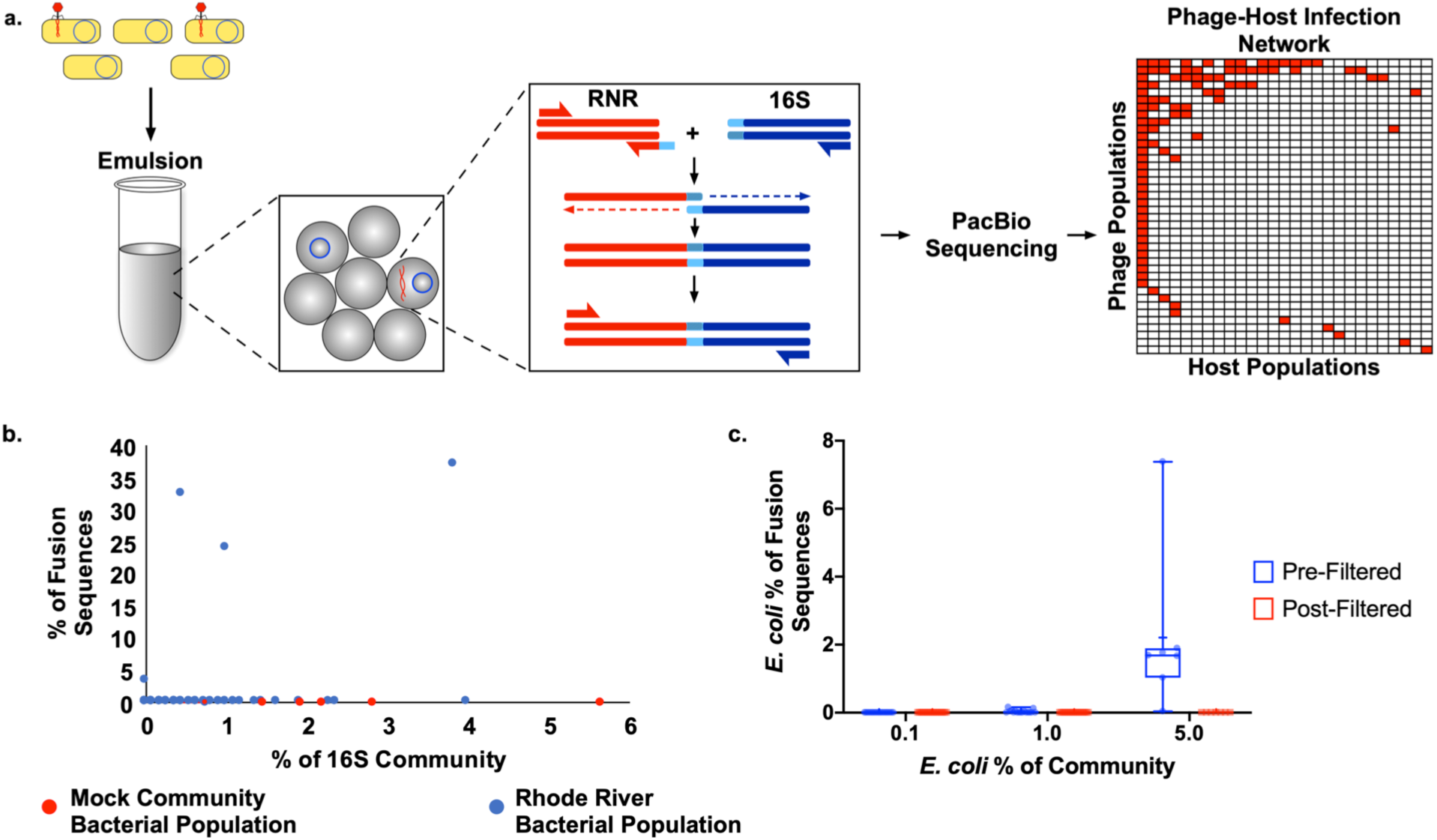
EpicPCR identifies phage-host interactions in the environment without cultivation. A) Overview of the experimental design for epicPCR, identifying phage-host interactions through single-cell isolation in emulsion droplets and virus and host marker gene fusion. Left: individual cells are isolated within emulsion droplets and the genome of the host (blue circle) and virus (red curved lines) serve as the template for the gene fusion reaction. Middle: fusion PCR joins and amplifies viral and host marker genes from actively infected cells within emulsion droplets. RNR (ribonucleotide reductase; red) and 16S (16S rRNA gene; blue) are joined through an overlapping primer sequence (light blue). Right: fused amplicons are sequenced and analyzed to identify the network of viral-host interactions in the environment. Overall, 95 interactions (red cells) between 40 unique phage RNR sequences and 27 unique host 16S rRNA sequences were identified. B-C) Specificity of the method was tested by spiking Rhode River water samples with a mock community or single uninfected host (*E.* coli) prior to emulsification. B) Proportion of fusion sequences belonging to uninfected mock (red) and Rhode River (blue) community members as a function of their community abundance. Uninfected mock community sequences were not found to be associated within any fusion products. C) Proportion of fusion products containing *E. coli* when spiked into Rhode River samples at 0.1% (n = 17), 1% (n = 17), and 5% (n = 7) of the community. Post-filtered sequences are those interactions observed in a minimum of three libraries.

We validated the specificity of epicPCR using uninfected mock communities and spike-in controls. With a complex mock community spiked into an environmental sample, uninfected control cells comprised as much as 16% of the total community and accounted for up to four of the ten most abundant community members. Despite their dominance, mock community control sequences were not observed in any fusion products (Fig. 3B) Additionally, we ensured specificity by adding uninfected *E. coli* cells to replicate epicPCR reactions from samples collected between May and December 2018, to control for non-specific interactions. Although uninfected control sequences were occasionally observed in the fusion data, these non-specific interactions were not consistently associated with any specific phage sequence. Thus, non-specific associations were removed by requiring that virus-host sequence pairs be observed in at least three libraries to be considered a positive interaction (Fig 3C).

EpicPCR analysis yielded 8,319 fusion amplicon sequences that shared 100% nucleotide identity across at least three samples (out of 58 total samples). The fusion amplicons contained 27 unique hosts (16S rRNA gene sequence at 100% identity), 40 unique phages (RNR at 100% identity), and 95 unique phage-host interactions between May and December 2018 (Fig. 4). The RNR genes identified from the epicPCR experiment formed three distinct phylogenetic clades (Fig. 4A) and shared greatest homology with T7-like Chesapeake Bay cyanopodoviral isolates S-CBP1, S-CBP3, and S-CBP4 but had less than 80% nucleotide identity to any reference sequence (Table S1). Recruiting viral metagenome reads to the RNR amplicons demonstrated these viral RNR sequences were not very abundant in our samples and highlights the sensitivity of the epicPCR method (Fig. S5). These RNR sequences failed to assemble into longer contigs, likely due to their low abundances and micro-diversity, which can prevent the assembly of even abundant phage genomes^40^.

**Figure 4.**
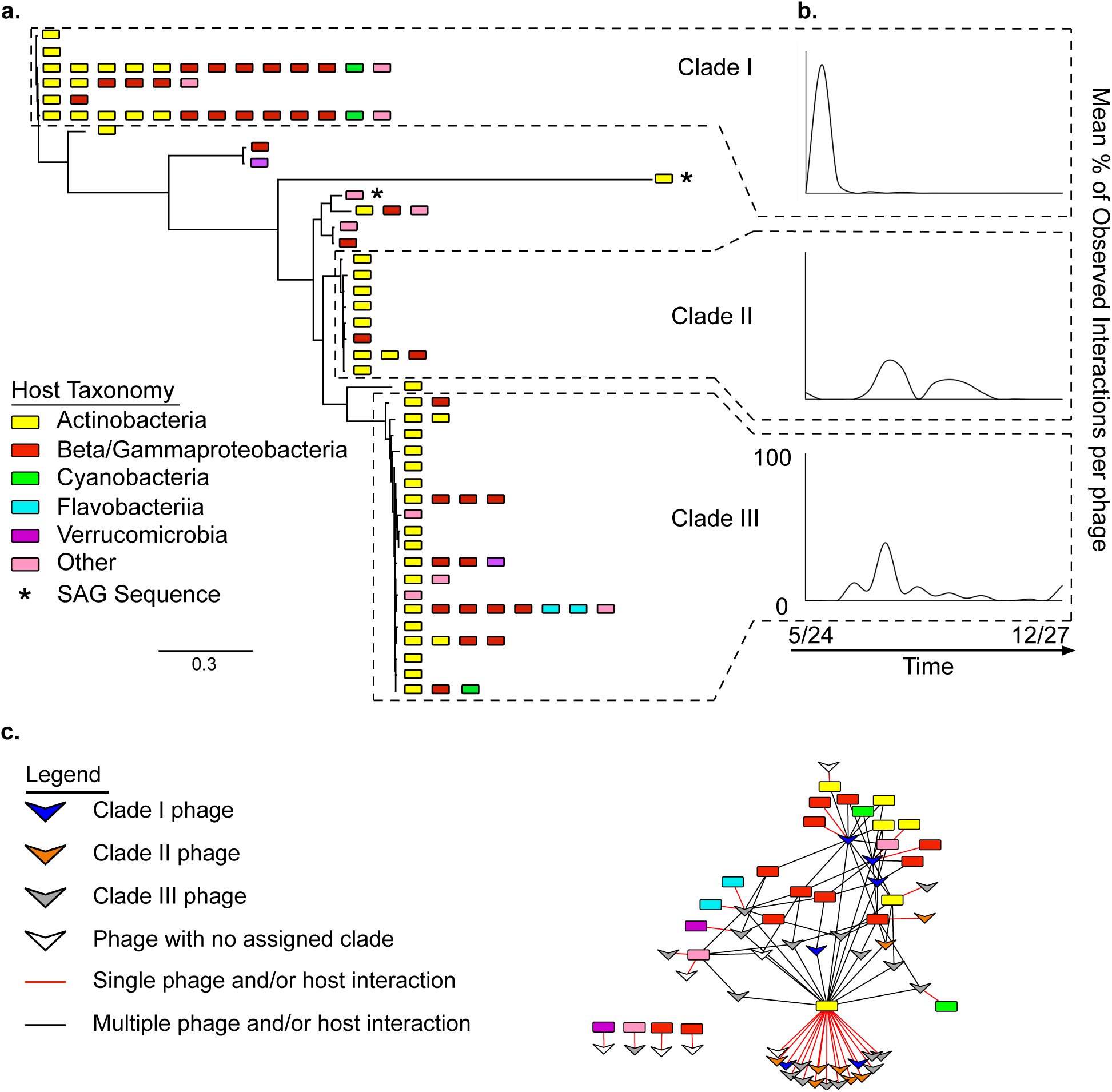
Abundance, diversity, and ecology of ‘Cyano SP’-like phage-host interactions. A) Maximum likelihood tree with 100 bootstrap replicates of Chesapeake Bay ‘Cyano SP’-like phage RNR gene sequences (G1380 to A2079 in *E. coli nrdA*). Only RNR gene sequences that were observed with the same host (100% nucleotide identity of 16S rRNA gene) in at least three epicPCR fusion amplicon libraries were included. Phage RNR sequences were clustered at 100% nucleotide identity and host representatives were plotted on the tree. Host taxonomy identified by 16S rRNA gene homology for each phage RNR gene sequence is indicated. Scale bar represents nucleotide substitutions per site. B) Mean percentage of total identified interactions observed at each sample time point by phage RNR clade. Clade I phage RNR sequences were primarily observed in late spring, while RNR sequences from Clades II and III were observed throughout the summer. C) Aggregate network of Chesapeake Bay ‘Cyano SP’-like phage-host interactions from May – December 2018. Host taxonomy is colored as in panel A. Most ‘Cyano SP’-like phage RNR sequences were associated with a single Actinobacteria host 16S rRNA gene identified as Luna-1 member *Rhodoluna*.

Unexpectedly, RNR sequences spanning all three clades were most frequently associated with Actinobacteria (Fig. 4). Approximately 80% (31/40) of the RNR sequences were linked to a single host classified as Actinobacteria Luna-1 subcluster member *Rhodoluna*, indicating this is the primary host for these particular Chesapeake cyanopodoviral-like populations. Unlike populations of Cyanobacteria (e.g. *Synechococcus* abundance varied > 1000-fold between July and December), this host displayed much less temporal variation in abundance (15-fold change between July and December), which may explain the observed stability of putative cyanopodoviral-like populations (1.1-fold change) compared to cyanomyoviruses (52-fold change) in virome libraries across the same time-points. Screening single-amplified genome (SAG) libraries further supports this observation. No Cyano SP-like RNR gene sequences could be amplified from ∼300 Cyanobacteria SAGs from May 2017. In contrast, one Cyano SP-like RNR gene sequence was associated with an Alphaproteobacteria SAG, while a second was associated with an Acidimicrobiia (phylum Actinobacteria) SAG library (Fig. 4A), providing further support that poorly characterized Chesapeake Bay viral populations with Cyano SP-like RNR genes interact with Actinobacteria.

Actinobacteria, including *Rhodoluna*, were also commonly predicted hosts of Chesapeake Bay viral populations using an *in silico* approach that infers the hosts through compositional similarity between host and viral genomes (WiSH^41^). The host genome database was created from references genomes^42^ and metagenome assembled genomes (MAGs) from the environmental bacterial fractions. To identify significant host predictions, the viral database included both viral populations and viral reference genomes. This resulted in significant (p < 0.05) host predictions for about half (49%) of the viral populations (Fig. S6). Most of the top host predictions for Chesapeake Bay viral contigs were reference genomes (80%); however, four of the ten most frequently predicted hosts were Chesapeake Bay MAGs, all of which were classified as Actinobacteria. In fact, among Chesapeake Bay MAGs Actinobacteria were the predicted hosts for more viral contigs (57%) than all other MAGs combined even though Actinobacteria only comprised 25% of MAG classifications overall (Fig. S6). Viral populations with putative Chesapeake Bay Actinobacteria hosts were abundant and consistently observed throughout the time series (Fig. S7), suggesting significant viral pressure on Actinobacteria populations throughout the year.

Incubation experiments of water samples with and without active viruses from the same site one year after the initial sample collection (July 2019) also suggest substantial viral pressure on Actinobacteria lineages (Fig. S8). Actinobacteria lineages Acidimicrobiia and Actinobacteria (class level) were the most commonly identified taxa susceptible to viruses, as determined by differences in relative abundance between incubations with active and inactive virus fractions (Fig. S8B). An identical 16S rRNA gene sequence to the primary *Rhodoluna* host identified by epicPCR was in the top half of the most susceptible populations, while a sequence with one nucleotide difference from the *Rhodoluna* host was in the top third. Previously, Actinobacteria was the most frequently identified host for a set of over 2,000 freshwater phage genomes in an analysis of freshwater viruses^43^, suggesting that Actinobacteria may be under substantial viral pressure in both freshwater and estuarine environments.

### In situ *viral-host interaction dynamics reveal ecologically differentiated viral clades*

EpicPCR analysis revealed interaction patterns that suggest RNR viral clades are ecologically differentiated. Phage-host interactions displayed seasonal patterns consistent with those observed for the viral community as a whole. The total number of interactions identified among Clade I ‘Cyano SP’-like RNR phage sequences peaked in late May before largely disappearing during the summer (Fig. 4B). Clade I phage sequences were most frequently associated with multiple hosts (4/6 sequences) and the greatest number of hosts (Fig. 4A, C). Host ranges for Clade I phages were significantly correlated with ‘Cyano SP-like’ phage abundances measured by qPCR (Spearman’s Rho = 0.69; p < 0.01). Thus, the broader host ranges of Clade I phages could be the temporary byproduct of higher viral abundances promoting less efficient interactions through increased contact rates. Cultivation studies have shown that bacterial hosts are not infected by different phages with equal efficiencies, as host sensitivity to phage infection can differ by orders of magnitude at the same phage titre^44^. It is also possible that broader host ranges provide an advantage for late spring viral populations under dynamic conditions characterized by large freshwater influxes.

In contrast, Clade II and III phage-host interactions were primarily observed during the summer (Fig. 4B). Most (7/8) Clade II phage sequences were associated with a single host, while clade III sequences were more evenly split between single hosts (11/19 sequences) and multiple hosts (8/19 sequences). Neither clade displayed the broad host ranges nor the correlation between host range and viral abundance observed with Clade I phage sequences. During this time, interactions between *Rhodoluna* and Clade II and Clade III phage populations alternated at most time points (Fig. 5), which may be a reflection of tidal influences on phage and/or host diversity. Supporting this hypothesis, Clade III interactions were negatively correlated with fluorescent dissolved organic matter (FDOM) (Spearman’s Rho = 0.67; p < 0.01), which can be strongly influenced by tide at our sample site^45^, and positively correlated with salinity (Spearman’s Rho = 0.57; p < 0.05) (Table S2). This suggests Clade II and III phage populations may represent riverine and estuarine diversity, respectively. However, we cannot rule out other possible explanations, such as changes to host physiology with different environmental conditions^46^ or antagonistic evolution between phage and host, as previously reviewed^47^.

**Figure 5.**
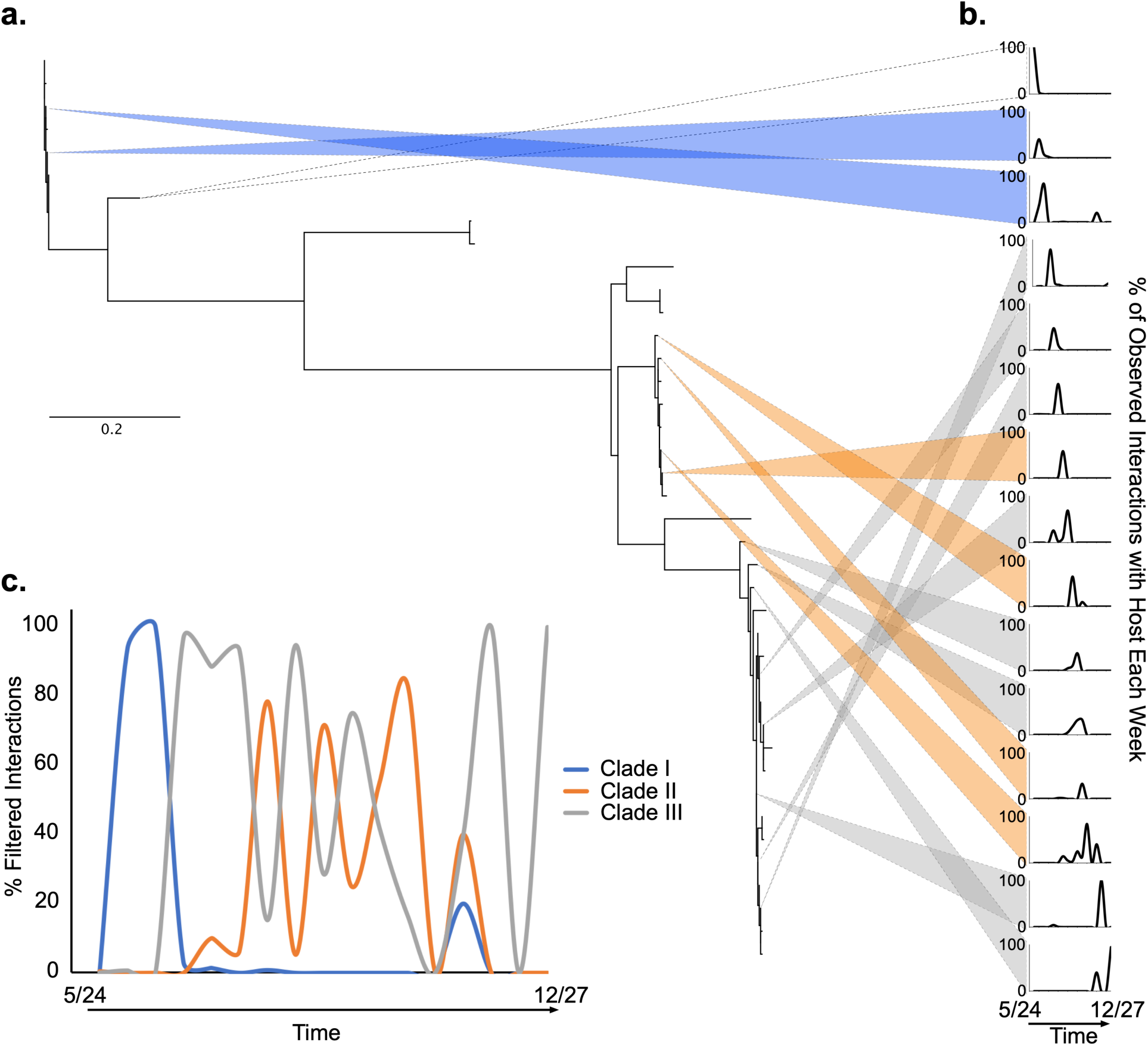
Phage interactions with *Rhodoluna* host observed from 5/24/18 to 12/27/18. In total, 31 of 40 phage populations interacted with this host. A) Maximum likelihood tree with 100 bootstrap replicates of Chesapeake Bay phage ribonucleotide reductase (RNR) genes (G1380 to A2079 in *E. coli nrdA*). RNR gene sequences were clustered at 100% nucleotide identity prior to phylogenetic analysis. Phage populations formed three main clades. Scale bar represents nucleotide substitutions per site. B) Proportion of total observed interactions with host that individual phage populations comprised each week. Colored wedges link phage populations to their corresponding dynamics over time. Wedge colors correspond to phage clades. Only the most abundant phage interactions were depicted for simplicity. C) Proportion of observed phage interactions with host over time by clade. Blue: Clade I; orange: Clade II; grey: Clade III.

### Host range associated with different virus-host interaction lifespan and efficiency

Interactions captured by epicPCR illuminated differences between viral RNR sequences associated with a single host (specialist phage) and multiple hosts (generalist phage) with regards to *in situ* interaction persistence and interaction efficiency. Generalist phage RNR sequences were observed in epicPCR fusion amplicons across a significantly (p < 0.01) greater number of sample dates than were specialist phage RNR sequences (Fig. 6A). This suggests that broader host ranges may increase the ‘interaction lifespan’ of phage and could be an evolutionary advantage in highly dynamic systems where viral infections appear to be largely ephemeral ^48,49^ (Fig. 4). However, epicPCR and 16S rRNA marker gene analyses also revealed differences in minimum host abundances when associated with generalist or specialist phages. Overall, 21 host sequences (100% nucleotide identity) that were associated with generalist phages from epicPCR were identified in the 16S rRNA libraries. The minimum abundance of these hosts when associated with a generalist phage RNR sequence ranged from 0 – 2.1% of the community (0.63% mean). Eight hosts associated with specialist phages from epicPCR were also identified in the 16S rRNA libraries and had minimum abundances ranging from 0 – 0.33% of the community (0.11% mean). Thus, the average minimum abundance of hosts when associated with specialist phage RNR sequences was significantly (p < 0.01) lower than when associated with generalist phage RNR sequences (Fig. 6B). This could be attributed to increased infection efficiencies among specialist phages compared to generalist phages. Consistent with these findings, previous work identified fitness costs that were associated with increased host range in cultivated phage-host systems^50^, and models of phage-host interactions suggest a trade-off in viral infection efficiency with increasing host range^12^. Considering the differences we observed in interaction persistence and interaction efficiency, there appears to be a trade-off between ‘interaction lifespan’ and infection efficiency for ‘Cyano SP’-like phages in the Chesapeake Bay.

**Figure 6.**
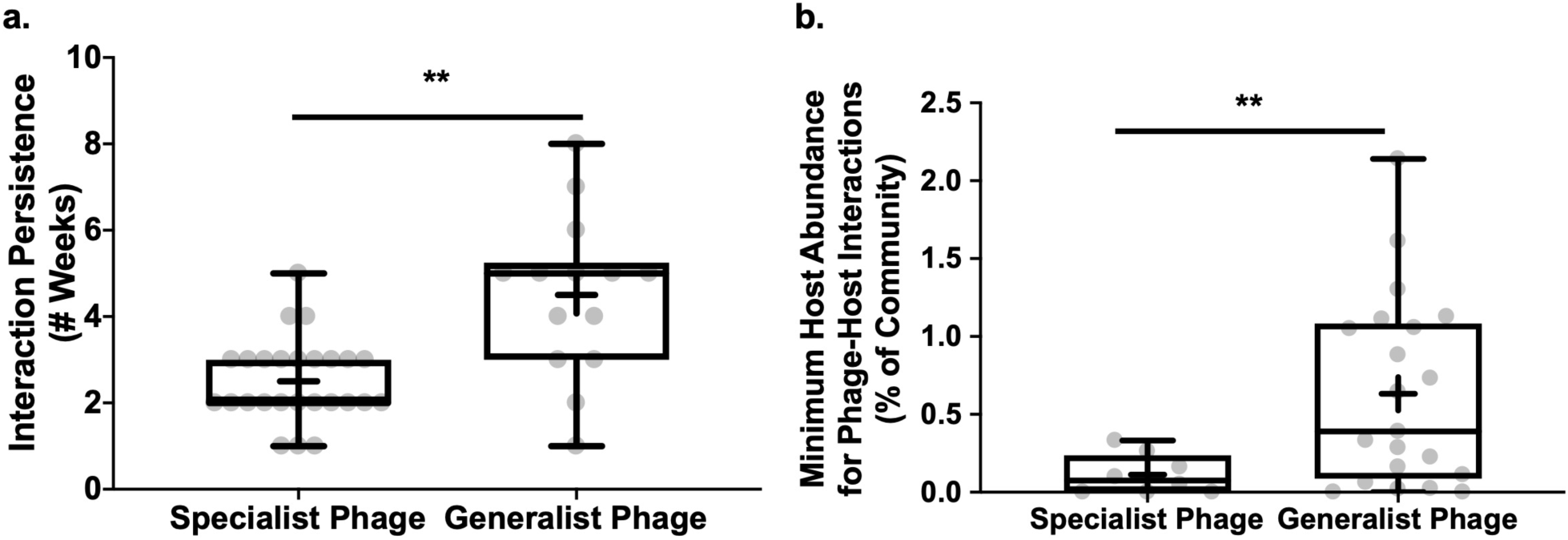
Biological trade-offs and ecological patterns related to viral and bacterial interactions. A) Total interaction persistence for ‘Cyano SP’-like phage RNR sequences associated with a single host (specialist phage, n = 26) or multiple hosts (generalist phage, n = 14) between May and December 2018 in the Chesapeake Bay (** = T Test p < 0.01). B) Minimum host abundance (16S rRNA gene relative abundance) of interactions with specialist phage RNR gene sequences (n = 8) and generalist phage RNR gene sequences (n = 21) observed in epicPCR fusion amplicons from samples collected between May and December 2018 in the Chesapeake Bay (** = T Test p < 0.01).

## Discussion

EpicPCR fills a gap as a culture independent, high-throughput tool that can complement existing methods in viral ecology by linking viral and bacterial populations *in situ*. EpicPCR can complement large-scale metagenomic sequencing efforts^51,52^ through host identification for novel and uncultivable viruses, a notable advantage over cultivation-based approaches, like high-throughput methods like viral tagging^53^. The method also has a throughput advantage over similar methods like digital PCR and single-cell genomics, which have also been applied to identify host interactions^22,54^ but can currently only accommodate hundreds to a few thousand wells for reactions and require costly and/or specialized equipment. In contrast, hundreds of thousands of reactions are possible within an emulsion with epicPCR, and the method can be performed using standard molecular biology equipment. Although primer bias remains an issue for epicPCR, the viral marker gene could be modified to any of the commonly applied viral marker genes^55^, or new primers could be developed to target novel viral diversity. For example, epicPCR could be used to examine host ranges of abundant viruses in the ocean, such as vSAG 37-F6 ^56,57^, or to verify specific host predictions, such as for potential archaeal viruses^58^. Recently, metagenomic time series have been used to predict environmental viral-host infection networks^59^. Observations from epicPCR could inform and test the accuracy and underlying assumptions of predicted network, improving our understanding of the drivers shaping microbial diversity and function. Unlike cultivation-based methods or *in silico* predictions, epicPCR can also capture virus-host interactions when and where they occur. This can be particularly useful for investigating how ecological and environmental factors influence these interactions. It should be noted, however, that while epicPCR captures close physical contact consistent with infections, it does not confirm active infections or differentiate between lytic or lysogenic infections, similar to the limitations of single-cell genomics^60^. However, pairing epicPCR with metatranscriptomics to examine expression of the viral marker gene^48^ could confirm active infections. Finally, pairing the host range capabilities of epicPCR with quantitative methods of viral abundance, such as qPCR^61^, ddPCR^57^ or polonies^62^, could be used to estimate the impact of viruses on different microbial taxa and facilitate estimates of the viral influence in ecosystem-level models.

In this study, we developed primers to target a relatively well-studied group of cyanophages and ascertain their *in situ* host interactions. This approach yielded unexpected host associations and interaction dynamics. Our analysis revealed that these ‘Cyano SP-like’ RNR populations commonly interact with Actinobacteria, which could not have been anticipated from homology searches of any phage genomes known to be associated with cultured Actinobacteria phage, such as within PhagesDB^63^. These associations are robust across methodologies, as evidenced by the identification of an ‘Cyano SP-like’ RNR sequence within the DNA amplified from a sorted, single cell with a 16S rRNA gene sequence that was classified as Actinobacteria. In addition, some of the phages that interacted with Actinobacteria were also associated with other hosts that spanned phyla. Although most viruses are believed to have narrow host ranges, host enrichment may select against broad-host range phages^64^ and cultivated viruses capable of infecting hosts spanning taxa at the order level have been reported^65^. Additionally, Paez-Espino et al. (2016)^66^ found one percent of viruses from metagenomic libraries had predicted hosts that spanned phyla based on CRISPR spacer and transfer RNA homology, including phages that were associated with Actinobacteria. EpicPCR may be better suited to capture fleeting or inefficient associations with alternative hosts than other methods like plaque assays, where infection and lysis by low-virulence phages can go undetected^67^, and could explain the greater frequency of inter-phyla host ranges in this data set. Furthermore, cultivation-based host range experiments often screen against related hosts. We most frequently observed generalist phages associated with *Rhodoluna* (Actinobacteria) and the Nor5-3 clade of Gammaproteobacteria, which would be unlikely to be screened together. It is also possible that these broad host ranges represent artifacts of the epicPCR method even though we used controls in all environmental samples to remove spurious interactions. However, observations of multiple different phages associated with the same hosts (*Rhodoluna* and Nor5-3 Gammaproteobacteria) suggests this is unlikely.

EpicPCR enabled us to identify differences in interaction persistence and interaction efficiency between generalist and specialist phage that would not have been identifiable in shotgun sequence data, which can be used to test specific hypotheses, such as the impact of generalist or specialist phages on community structure or the relative contribution of specific interactions to viral production and bacterial mortality. Since generalist predators tend to have a stabilizing effect on the overall community diversity at both the macro^68^ and micro scale^69^, we would hypothesize that when Clade I infections are dominant, such as in the spring, they would tend to stabilize community alpha diversity, while when Clade II infections are dominant, such as with increasing amounts of FDOM, they would tend to destabilize diversity. These interactions would be one in a myriad of interactions that would shape the overall community diversity, but if these specific interactions had a disproportionate impact on the community-level processes, the stabilizing and destabilizing effects might be observable at the community level. Virus-host interaction dynamics revealed by epicPCR shed light on the individual interactions that contribute to viral production and bacterial mortality over time and with changing conditions. Previous studies revealed differences in viral productivity and bacterial mortality associated with spring and neap tides^8^. While our study did not quantify viral productivity or bacterial mortality, we did find that interactions between *Rhodoluna* and Clade II and Clade III phages were associated with salinity changes. Advances to this technique, such as pairing with a microfluidics-based droplet maker and sorting and uniquely barcoding positively amplified droplets may yield quantitative results, although these advances would limit wide-spread accessibility. In this way, such quantitative measures could be paired with measurements of viral productivity and bacterial mortality to determine the relative contribution of specific interactions to viral production and bacterial mortality.

## Conclusions

Viruses influence coastal ocean and estuarine biogeochemistry through interactions with hosts mediating key autotrophic and heterotrophic processes. Using a combination of shotgun metagenomics, marker gene analysis, dilution experiments and a novel gene fusion technique, we investigated the factors that influence virus-host interactions within the largest estuary in the United States. This estuary has a unique and endemic viral community that varies more between seasons than between years. We found Actinobacteria populations are under substantial viral pressure within this ecosystem and their interactions with specific phage clades seem to vary with tidally-influenced environmental factors. The observations of virus-host interactions from epicPCR also suggest that single- and multi-host phages have substantial differences in their ‘interaction lifespan’ and realized host abundance characteristics. This combination of bioinformatics and experimental approaches provided high genetic and temporal resolution of viral interactions with one of the most abundant heterotrophic bacterial populations within this ecosystem that can be widely applied across aquatic ecosystems to gain insight into viral ecology.

## Materials and Methods

### Sample Collection and preservation

Surface water samples were collected from the mouth of the Rhode River (Edgewater, MD) off of the Smithsonian Environmental Research Center (SERC) research pier. Samples were collected five times a day over three days from 05/17/17-05/19/17, between 10:30 am and 4:30 pm. Samples were also collected at 12:00 pm weekly from 5/24/18 to 8/9/18, on 8/23/18, then again weekly from 12/6/18 to 12/27/18. Briefly, 25 mL of water sample was combined with 25 mL of 50% (v/v) sterile glycerol. Glycerol samples were stored on dry ice for transport back to Baltimore, MD and subsequently stored at -80°C until processing. Approximately 120 mL of water sample was also collected per time point and filtered through a 0.2 µm PES membrane filter (Millipore, Inc.) for shotgun metagenomic analyses. Filters were stored on ice for transport back to Baltimore, MD and kept at -80°C until processing. Viral filtrates (< 0.2 µm fraction) were collected and incubated with FeCl_3_ as previously described ^70^ during transport back to Baltimore, MD. Viral filtrates were filtered through a 0.2 µm PES membrane filter (Millipore, Inc.) post-incubation, and filters were stored in the dark at 4°C until processing for shotgun sequencing. Water conditions were recorded from the continuous water monitoring station located at SERC (http://nmnhmp.riocean.com, Table S3).

### Bacterial and viral gene counts

Bacterial and viral abundances were estimated by quantitative PCR. To create a standard curve of gene copy number, viral ‘Cyano SP’-like RNR genes were amplified from environmental samples using primers Cyano_II_F and Cyano_II_R (0.3 uM final concentration each, Table S4). Amplicons were run on a 1.5% agarose gel and visualized with SYBR Safe DNA gel stain (Invitrogen, 1x final concentration). RNR bands were cut out, gel purified (Zymo, Inc.), and cloned into chemically competent *Escherichia coli* cells using the Zero Blunt PCR Cloning Kit (Thermo Scientific) following the manufacturer’s protocol. Cells were grown overnight on LB + kanamycin (50 µg mL^-1^) plates at 37°C and colonies were picked for subsequent testing. Picked colonies were tested for amplification of the RNR gene. One colony was serially diluted and grown on plates to determine gene copy number per dilution. The serial dilution series was used to create a standard curve for use in qPCR. All standards and environmental samples were run in triplicate. Three microliters of sample were combined with UltraPure molecular grade water (Thermo, Inc.), SsoAdvanced Universal SYBR Green Supermix (1x final concentration, Bio-Rad Laboratories, Inc.), Cyano_II_F primer (0.3 μM final concentration), and Cyano_II_R primer (0.3 μM final concentration) to a final volume of 25 μL (see Table S4 for primer sequences and citations). Samples were amplified on a CFX96 Real-Time PCR Detection System (Bio-Rad Laboratories, Inc.) with the following conditions: denaturing at 98°C for 10 minutes; 45 cycles of denaturing at 98°C for 10 seconds, annealing at 52°C for 30 seconds, and extension at 72°C for 45 seconds; and a final extension of 72°C for 5 minutes. The same dilution series was used to create a standard curve for the 16S rRNA gene. 16S rRNA gene counts were assessed as above with primers PE_16S_U515F and 16S_1114R (Table S4). Bacterial cell counts in environmental samples were predicted based on the gene counts and copy number correction between *E. coli* (7 copies) and aquatic bacteria (approximately 3 copies).

### Shotgun metagenomic library preparation, sequencing, and processing

DNA was extracted from filters for shotgun sequencing from water samples collected on the following dates: 05/17/17, 05/18/17, 05/19/17, 05/31/2018, 06/28/18, 08/02/18, and 12/6/18. Additionally, two positive controls were processed, a Zymo positive control community (Zymo Research) and *E. coli*, and one negative control (water). DNA extraction was performed with the DNeasy PowerWater kit (Qiagen) following the manufacturer’s protocol with the following amendment: 20 µL of proteinase K was combined with 1 mL of solution PW1 in the bead tube. The bead tube was incubated at 65°C for ten minutes prior to bead beating. Libraries were prepared with the Nextera DNA Flex Library Prep kit (Illumina, Inc.) following the manufacture’s protocol and sequenced on an Illumina MiSeq (2 x 300 bp) at the Genetic Core Research Facility at Johns Hopkins University.

Sequences were quality filtered and trimmed with trimmomatic (v. 0.38)^71^, assembled with metaSPAdes (v. 3.13.1)^72^. The assembly was used to generate metagenome assembled genomes (MAGs) through metaWRAP^73^ with metabat^74^ and maxbin2^75^. Binning_refiner^76^ was used to create the final set of MAGs with at least 50% completeness and less than 10% contamination, as determined by CheckM^77^. MAGs were classified using Kaiju^78^

### 16S rRNA gene amplicon library preparation, sequencing, and processing

16S rRNA genes were amplified from 2017 and 2018 Chesapeake Bay surface water samples (see Fig. S1) in a 25 μL PCR reaction with the following conditions: three microliters of column-purified DNA were combined with UltraPure molecular grade water (Thermo, Inc.), 10X buffer (1x final concentration), dNTPs (0.1mM each final concentration), 16S forward primer 27F (0.3 μM final concentration), 16S reverse primer PE_16S_V4_E786_R (0.3 μM final concentration), bovine serum albumin (0.02 mg/mL final concentration), and Phusion High-Fidelity DNA Polymerase (0.5U; New England BioLabs, Inc.; see Table S4 for primer sequences and citations). PCR reactions were combined with 150 μL of 4% UMIL EM90 oil (4% UMIL EM90 oil, 0.05% TritonX-100 v/v in mineral oil; Universal Preserv-A-Chem, Inc.) and emulsified by vortexing at max speed (∼2,700 rpm) for one minute on a Vortex Genie 2 (MoBio). Emulsions were loaded as 50 μL aliquots and amplified with the following conditions: denaturation at 94°C for 3 minutes; 33 cycles of denaturation at 94°C for 10 seconds, annealing at 54°C for 30 seconds, and extension at 72°C for 45 seconds; and a final extension of 72°C for 5 minutes (C1000, BioRad Labs., Inc.). Samples were immediately removed upon completion of amplification and stored at -20°C until the emulsion was broken.

PCR oil emulsions were broken with isobutanol as previously described^79^. Briefly, PCR aliquots were pooled in a 1.5mL microcentrifuge tube and combined with 100 uL of sterile 5M NaCl solution and 1 mL of isobutanol. Samples were vortexed briefly to mix and centrifuged at 16,000 x g for 1 minute. The bottom aqueous layer was retained, and DNA was purified by spin column purification (Zymo, Inc.). DNA was eluted in 20 uL of Tris-HCl and stored at -20°C.

Purified DNA was run on a 1.5% agarose gel (UltraPure Agarose, ThermoFisher Scientific) and visualized with SYBR Safe DNA gel stain (Invitrogen, 1x final concentration). The gel was run in 1X TBE buffer (Alfa Aesar) at 4 V/cm. 16S rRNA gene bands were visualized under blue light excitation, extracted, and gel purified (Zymo, Inc.) Purified DNA was eluted into 20 μL of Tris-HCl and stored at -20°C until further processing.

Barcodes and Illumina adapters were added to 16S rRNA gene amplicon products in two subsequent limited PCR steps. Barcodes were added as follows: two microliters of purified DNA were combined with UltraPure molecular grade water (Thermo, Inc.), 10X buffer (1x final concentration), dNTPs (0.1mM each final concentration), 16S forward primer PE_16S_V4_U515F (0.3 μM final concentration), 16S rRNA gene reverse primer with 8-mer barcodes PE_IV_XXX (0.3 μM final concentration), and Phusion High-Fidelity DNA Polymerase (0.5U; New England BioLabs, Inc.; see Table S4 for primer sequences and citations). Samples were amplified with the following conditions: denaturing at 98°C for 30 seconds; 8 cycles of denaturing at 98°C for 10 seconds, annealing at 54°C for 30 seconds, and extension at 72°C for 45 seconds; and a final extension of 72°C for 5 minutes. DNA was purified by spin column purification (Zymo, Inc.) and eluted into 20 μL Tris-HCl. Illumina adapters were then added as above with the following primers: Illumina adapter forward primer PE-III-PCR-F (0.3 μM final concentration) and Illumina adapter reverse primer Barcode_Rev (0.3 μM final concentration) (see Table S4 for primer sequences and citations). Samples were amplified with the following conditions: denaturing at 98°C for 30 seconds; 5 cycles of denaturing at 98°C for 10 seconds, annealing at 54°C for 30 seconds, and extension at 72°C for 45 seconds; and a final extension of 72°C for 5 minutes. DNA was purified by spin column purification (Zymo, Inc.) and eluted into 20 μL Tris-HCl.

16S rRNA gene amplicon products were quantitated on a Qubit 3.0 fluorometer (Invitrogen) and three nanograms of DNA pooled per sample for sequencing. 16S rRNA gene amplicon libraries were sequenced on an Illumina MiSeq (2 x 300 bp) at the Genetic Core Research Facility at Johns Hopkins University. Sequence reads were processed in QIIME2^80^ using the DADA2 de-noising pipeline. Taxonomic assignment was performed in QIIME2 with the Greengenes^81^ database.

### Virome shotgun metagenomic library preparation, sequencing, and processing

Samples from the following dates were used for virome short-read shotgun metagenomic analysis: 05/17/17, 05/18/17, 05/31/18, 08/02/18, 12/14/18, 12/20/18. The two samples collected on 12/14/18, 12/20/18 were also long-read sequenced. Viruses were resuspended from filters with an ascorbic acid buffer as previously described^70^. Following resuspension, viral particles were purified by cesium chloride gradient centrifugation ^82^ and DNA extracted with Wizard Prep Columns (Promega, Corp.). Viral metagenome short-read libraries were prepared using the NexteraXT kit (Illumina, Inc.) following the manufacturer’s protocol. For samples with > 0.16 ng/μL, samples were amplified with 15 cycles; for those with 0.1-016 ng/μL, samples were diluted 1:5 and amplified with 18 cycles; for <0.1ng/μL, samples were diluted 1:10 and amplified with 20 cycles. Short reads for all virome libraries were sequenced on an Illumina Short reads for all virome libraries were sequenced on an Illumina NovaSeq S4 with 75M (target) 2×150bp reads at the JP Sulzberger Genome Center (Columbia University, New York, NY). Additionally, the long-read libraries from December 2018 were prepared using phenol:chloroform extraction protocol (dx.doi.org/10.17504/protocols.io.6cbhasn) and libraries were prepared as previously described ^24^ with modifications and sequenced with an Oxford Nanopore MinION instrument on a FLO-MIN106D R9 version Spot-ON flowcell (Rev D) at the Ohio State University according to the manufacturer’s instructions.

Short reads were cleaned and quality-trimmed with bbduk ^83^; adapters, sequencing artifacts, and PhiX sequences were removed (ktrim=r; k=23 mink=11; hdist=1; hdist2=1). Reads were then quality-trimmed from both ends to remove bases with low quality scores (qtrim=rl; trimq=20). Reads shorter than 30 bp (minlength=30), with Ns (maxns=0), or with an average quality below 20 (maq=20) were discarded. The cleaned reads from each sample were then independently assembled with metaSPAdes (v. 3.13.1)^72^ using k-mer sizes: 21, 33, 55, 77, and – meta parameter. Long-reads were basecalled with Guppy v.2.3.1 (Manufacturer’s tool) and individual barcoded sample libraries were demultiplexed with the ‘barcoder’ function of Guppy. Long-reads were quality controlled with NanoFilt ^84^, in which reads were filtered by quality score (Q-score ≥ 10) and minimum length (≥1kb), and finally ‘headcropped’ by 50bp to ensure no remaining barcode sequence remained. These cleaned long-reads were used in two separate assembly scenarios. First, long-reads were assembled with Flye ^85^ in metagenomic mode (--meta). Error-correction of the Flye assemblies where performed with Pilon v.1.23 ^86^, using the corresponding short-reads mapping information [read recruitment was performed with BWA v.0.7.17 ^87^] to detect and reduce basecalling errors. Second, hybrid assembly, using both long- and short-reads was performed with SPAdes (v.3.13.1) ^72^. Long reads were assembled through hybrid assembly (with metaSpades hybrid option) and Flye ^85^. In order to predict viral contigs in the assembled datasets, assemblies were run through VirSorter ^88^ (v. 1.0.5) in the -virome mode after upgrading its database with an expanded profile hmm database of viral proteins, mainly from the GOV2.0 dataset^14^. Contigs that were resolved as categories 1, 2, 4 and 5 were retained, and filtered by length ≥ 5kb (≥1.5kb, if circular). DeepVirFinder ^89^ was another tool for rescuing additional viral contigs. We considered high-confidence viral contigs from DeepVirFinder to be those of scores of ≥ 0.9 with a p-value ≤ 0.05, and lengths ≥ 5kb. These two sets of predicted viral contigs (from both long- and short-read assemblies) were then dereplicated using “ClusterGenomes” ^90^ into viral populations using 95% average nucleotide identity over 80% coverage of the shorter contig length.

To calculate the coverages of these viral populations, clean reads were mapped to the Chesapeake Bay viral population database with Bowtie2 ^91^ in the non-deterministic and sensitive mode. The output bam files were parsed using BamM (https://github.com/Ecogenomics/BamM) to only keep the reads that covered over 70% of the viral contig length, with over 75% read alignment length. Pysam (https://github.com/pysam-developers/pysam) was then used to filter out reads with <95% identity. Trimmed pileup coverage “tpmean” for each contig was calculated using BamM and then they were adjusted for each sample by metagenome size. The same read mapping strategy was employed for RNR sequences upon constructing the rank abundance curves in (Fig. S10)

To study seasonal diversity of Chesapeake Bay viral populations, all viromes were randomly subsampled to 15M reads using bbmap “reformat” ^83^ with default parameters. The subsampled read libraries were assembled using metaSPAdes (v. 3.13.1) as above. The resulting assemblies were then processed with VirSorter and DeepVirFinder as above. Viral contigs extracted using the same cutoffs as mentioned before were grouped into viral populations if they shared ≥ 95% nucleotide identity across ≥ 80% of the shorter contig length. Subsequently, the subsampled reads were recruited to the viral populations’ representatives with Bowtie2, and the same read-mapping cutoffs discussed above were applied to calculate sequence-depth adjusted coverage.Taxonomic assignment of viral populations was performed using vConTACT2 ^90,92^ with default parameters. First, the full proteome of every viral population representative was predicted using prodigal^93^. The protein set was then combined with all the proteins from the phage and archaeal viruses in the NCBI RefSeq v88 release. The combined set of proteins was then used as input for vConTACT2 to compute protein similarity overlaps (i.e. protein clusters), and subsequently refined into genus-level equivalent ‘viral clusters’ (VCs). Using this method, viruses are classified at the genus level. The resulting cluster file were imported and visualized in Cytoscape 3.7.2^94^.

### Seasonal dynamics and diversity analyses

Seasonal bacterial communities were assessed from 16S rRNA gene amplicon libraries. Hierarchical clustering was performed in QIIME^95^ with Bray-Curtis dissimilarity. Libraries were sub-sampled at 150,000 observations with ten jackknifed replicates. Community structure was visualized using heatmap.2 in the gplots^96^ package in R.

Seasonal Chesapeake Bay viral communities were characterized by mapping reads to assembled contigs using Bowtie2^97^ with default sensitive conditions. To compare viral populations with a deeply sequenced virome library collected in 2012 at the same location^36^, 2017 and 2018 Chesapeake Bay virome reads were mapped to both the > 5 kb contigs assembled in this study and contigs > 5kb from the 2012 virome that were previously assembled. Only reads that mapped to contigs with ≥ 90% identity over ≥ 90% of the read were retained. Viral populations were considered present in the sample if retained reads mapped to ≥ 75% of the contig. Trimmed pileup coverage values were used as proxies of observation counts for each population. Hierarchical clustering was performed in QIIME with Bray-Curtis dissimilarity. Libraries were sub-sampled at 300,000 observations with ten jackknifed replicates. Community structure was visualized in R as described above.

Shannon diversity was calculated in QIIME for libraries with corresponding virome samples (5/17/2017, 5/31/2018, 8/2/2018, 12/14/2018, 12/20/2018). Values were recorded as the average of ten jackknifed replicates at a sampling depth of 10,000 observations for both 16S and viral populations.

The distribution of Chesapeake Bay viral populations in ocean viral metagenomes was investigated by mapping viral metagenome reads from Global Ocean Virome 2.0 database libraries^14^ to the Chesapeake Bay > 5kb contigs as described above for the Chesapeake Bay 2012 viral metagenome sample. Additional libraries from previously published freshwater and estuarine environments were similarly queried^29-31^, in addition to a viral metagenome from the Damariscotta River Estuary, ME, USA (BioProject Accession No. PRJNA357591).

### Ribonucleotide reductase homology analysis

Open reading frames (ORFs) were called for all Chesapeake Bay > 5kb contigs using MetaGeneMark^98^. Ribonucleotide reductase (RNR) alpha subunit genes were identified by BLASTx query of ORFs (e-value 1E-10) against a database of Uniref50^99^ RNR cluster representative sequences. Putative RNR genes were translated to amino acid sequences and aligned with MAFFT^100^. Aligned sequences were visually inspected in Geneious^101^ v. 9.1.5 for the presence of conserved catalytic residues C439, E441, C462 indicative of RNR proteins (amino acid positions of *Escherichia coli* nrdA gene product). Only sequences containing these conserved amino acids and spanning the C462 to P621were retained. All retained sequences were queried against viral sequences in the NCBI nr database by BLASTn to identify putative viral and host taxonomy. Top hits with a percent identity ≥ 65% across at least 90% of the query sequence were recorded. All sequences with top hits below this threshold were reported as ‘Unclassified’.

### Bioinformatics host prediction with WIsH

WIsH^42^ was used to determine potential hosts for viral contigs greater than 10 KB in length. The entire dataset used in the WIsH analysis included viral contigs and metagenome assembled genomes (MAGs >50% completion, <10% contamination) from the Rhode River, along with the viral and hosts reference genomes from the WIsH benchmark dataset, downloaded from NCBI. Because an appropriate null model is not available for this unique estuarine environment, we assume that for every bacterial model, the set of phage genomes in the dataset for which it is a host is negligible compared to the set of phage genomes that for which it is *not* a host. Reference host and viral genome relationships in the benchmark dataset were used to verify the performance under these assumptions. Running the benchmark dataset with the null model yielded similar results as those previously reported (58%, as compared to 63%, accuracy reported at the genus level). Thus, we applied the same assumption for the null model to the analysis of environmental data. The analysis required top hits to have a likelihood value in the top 5% of all calculated likelihoods (p <= 0.05). Of the 9,392 viral contigs from the Chesapeake Bay, almost half had top predictions that were within the top 5% of all tested likelihoods. Of significant hits, 80% of top predictions of hosts for environmental viral population mapped to reference genomes and 20% of significant top predicted hosts were MAGs.

### epicPCR of environmental samples

Glycerol samples were thawed on ice and one mL was added to three replicate 1.5 mL microcentrifuge tubes per sample. One replicate was left unamended, while the other two were spiked with *E. coli* to approximately 0.1% and 1% of the bacterial community, respectively, to identify false-positive interactions. A fourth replicate with 5% *E. coli* was processed for seven of the time points. Samples were centrifuged at 25,000 x g for 10 minutes and resuspended after supernatant removal to reduce free viral particles. Thirty microliters of each sample was combined with UltraPure molecular grade water (Thermo, Inc.), 10X buffer (1x final concentration), dNTPs (0.1mM each final concentration), Cyano_SP_F primer (1.0 μM final concentration), Cyano_SP_R_519R primers (0.01 μM each final concentration), S-*-Univ-1100-a-A-15 16S reverse primer (1.0 μM final concentration), bovine serum albumin (0.02 mg/mL final concentration), Tween-20 (0.8% v/v final concentration), and Phusion High-Fidelity DNA Polymerase (1.5U; New England BioLabs, Inc.) to a final volume of 75 μL (see Table S4 for primer sequences and citations). PCR reactions were combined with 450 μL of 4% UMIL EM90 oil (4% UMIL EM90 oil, 0.05% TritonX-100 v/v in mineral oil; Universal Preserv-A-Chem, Inc.) and emulsified by vortexing at max speed (∼2,700 rpm) for one minute. Emulsions were loaded as 50 μL aliquots and amplified with the following conditions: denaturation at 94°C for 3 minutes; 33 cycles of denaturation at 94°C for 10 seconds, annealing at 54°C for 30 seconds, and extension at 72°C for 45 seconds; and a final extension of 72°C for 5 minutes (C1000, BioRad Labs., Inc.). Samples were immediately removed upon completion of amplification and stored at -20°C until the emulsion was broken as described above.

### Preventing fusion of unfused genes during nested PCR

Blocking primers were used during nested PCR to prevent the annealing and amplification of unfused genes from the emulsion PCR as previously described^23^. The efficacy of the blocking primers was tested to ensure random virus-host sequences were not being introduced. First, RNR and 16S rRNA gene copy numbers were quantitated in all epicPCR reactions post-cleanup by qPCR as described above. Next, RNR genes were amplified from Rhode River samples with primers Cyano_II_F and Cyano_II_R_519R primers (0.3 μM final concentration each, Table S4). 16S rRNA genes were similarly amplified with primers 27F and 1492R (0.3 μM final concentration each, Table S4). Unfused RNR and 16S rRNA gene amplicons were combined at copy numbers equal to the highest observed across all samples after epicPCR as determined by qPCR. Finally, the unfused RNR and 16S rRNA gene mixture was run through nested PCR and barcoding PCR reactions with all environmental samples as described below. Samples were run on a 1.5% agarose gel and visualized with SYBR Safe DNA gel stain (Invitrogen, 1x final concentration). Fusion products were observed in the unfused mix control when no blocking primers were used. However, with the addition of blocking primers no fusion products were detected in the unfused mix control.

### Enrichment of viral-host fused amplicons

Fused amplicons were enriched by nested PCR. Three microliters of column-purified DNA were combined with UltraPure molecular grade water (Thermo, Inc.), 10X buffer (1x final concentration), dNTPs (0.1mM each final concentration), Cyano_SP_Nested_FA primer (0.3 μM final concentration), 16S reverse primer PE_16S_V4_E786_R (0.3 μM final concentration), forward and reverse blocking primers U519F-block10 and U519R-block10 (1.0 μM final concentration each; to ensure no amplification of unfused genes, see above), and Phusion High-Fidelity DNA Polymerase (0.5U; New England BioLabs, Inc.) to a final volume of 25 μL (see Table S4 for primer sequences and citations). Samples were amplified with the following conditions: denaturing at 94°C for 30 seconds; 30 cycles of denaturing at 94°C for 10 seconds, annealing at 54°C for 30 seconds, and extension at 72°C for 45 seconds; and a final extension of 72°C for 5 minutes. PCR reactions were cleaned by spin column purification (Zymo, Inc.) and DNA eluted in 20 uL of Tris-HCl. Samples were barcoded following enrichment and cleanup by PCR as described above with Cyano_SP_Nested_FB primer (0.3 μM final concentration, Table S4), reverse primer PE-IV-PCR-XXX with 8-mer barcodes (0.3 μM final concentration, Table S4), and forward and reverse blocking primers U519F-block10 and U519R-block10 (1.0 μM final concentration each, Table S4). Barcoded samples were run on a 1.5% agarose gel (Invitrogen) and visualized with SYBR Safe DNA gel stain (Invitrogen, 1x final concentration). Fusion bands were cut out and gel purified (Zymo, Inc.). Fusion products were quantitated on a Qubit 3.0 fluorometer (Invitrogen) and ten nanograms of DNA pooled per sample for sequencing.

### Sequencing and quality filtering of viral-host fusion amplicons

Fused amplicon products were sequenced on a PacBio Sequel with Sequel v3 chemistry (University of Maryland Institute for Genome Sciences). Circular consensus sequences were obtained from raw reads with the following parameters: minimum signal-to-noise ratio (SNR): 3, minimum length: 500bp, minimum passes: 10, minimum read score: 0.75, minimum predicted accuracy: 0.90. Consensus sequences passing these thresholds were oriented by searching for primer sequences using Cutadapt^102^ with an error rate of 0.01. Only sequences passing this error rate were retained. Following orientation, reads were demultiplexed in QIIME^95^ with zero allowed barcode mismatches. Fused gene products were split by gene and primer sequences identified with Cutadapt^102^. Only those sequences without mismatches in any primer sites (error rate of 0.01) were retained for further analyses. Finally, amplicons were filtered by size to remove truncated or chimeric reads. Only fusion sequences with RNR gene amplicons between 583 and 596bp and 16S rRNA gene amplicons between 249 and 262bp were retained. These sequences were split into separate RNR and 16S genes and their primer sequences trimmed with Cutadapt^102^.

### Analyzing viral and bacterial diversity from epicPCR fusion amplicons

Viral RNR and bacterial 16S rRNA gene sequences passing all quality filtering were clustered at 100% nucleotide identity with CD-HIT^103^ to identify viral and bacterial populations. Only viral-host pairs observed in three or more libraries were considered positive interactions. Representative sequences from these viral and bacterial populations were queried against the NCBI nr database via BLASTn to identify top hits. RNR gene sequences were aligned with MAFFT^100^ using the L-INS-I setting. Sequences were trimmed to the region G1380 to A2079 in *E. coli nrdA* and a maximum likelihood tree with 100 bootstrap replicates was made using Phylogeny.fr (http://phylogeny.lirmm.fr)^104^. Viral-host interaction networks were visualized in Cytoscape.

### Viral dilution incubation experiments

One liter of surface water was collected in July 2019 from the mouth of the Rhode River (Edgewater, MD) at the Smithsonian Environmental Research Center (SERC). 120mL was filtered through two 0.2 μm PES membrane filter (Millipore, Inc.) and the viral filtrate was collected. Half of the viral filtrate was autoclaved to eliminate active viruses. The remaining viral filtrate was left unamended. Bacterial communities were resuspended off of the 0.2 μm filters in 1mL autoclaved viral filtrate by shaking for 10 minutes at medium speed on a MoBio Vortex Genie 2. Resuspensions from each filter were pooled to create a single resuspended sample. Resuspended cells were combined 1:120 with either autoclaved (no viruses) or un-autoclaved (with viruses) viral filtrate. Samples were divided into twelve 20 mL replicates. Six replicates without viruses and six replicates with viruses were incubated at room temperature. 500 μL aliquots were taken at the beginning of the incubation and approximately every 24 hours over three days. 16S rRNA and ‘Cyano SP’-like RNR gene counts were quantified at each time point by qPCR as described above. 16S amplicon libraries were created as described above for the initial community and the community after 75 hours for all replicates. Growth of individual bacterial populations over the course of the incubation was calculated from qPCR results and their relative abundance. Only bacterial populations that grew in the absence of viruses were retained for further analysis. Susceptibility to viral-mediated mortality was calculated per bacterial population as (Population Abundance Fold-Change Without viruses)/(Population Abundance Fold-Change With Viruses). Higher values indicate greater susceptibility to viral-induced mortality. Growth rates were calculated per 16S bacterial population as the fold-change in abundance over time based on relative abundance and qPCR results. Relative growth rates were calculated per 16S bacterial population to normalize between replicates. Relative growth rate was defined as the growth rate of a bacterial population relative to the growth rate of the entire community during incubation without viruses.

### Single-cell genomics

Samples from May 18, 2017 were sent to the Bigelow Single Cell Genomics Center (East Boothbay, ME) for sorting of the cyanobacteria and prokaryotic fraction into 384-well plates. Sorting was conducted with the red fluorescence as a function of forward scatter, used to gate for “cyanobacteria”, and Syto-9 stained DNA as a function of forward scatter, used to gate for “all prokaryotes”. Single cell genome amplification (SAG Generation 2) of sorted cells was conducted at the Bigelow labs and returned to JHU for marker analysis. RNR and 16S rRNA marker genes were amplified from 2 μL of a 1:100 dilution of the amplified genomes according to the protocol above, without the use of emulsions. Positive amplicons were purified and sequenced with the forward primers at the Genetic Resource Core Facility at JHU with an Applied Biosystems 3730xl DNA Analyzer.

## Data availability

Sequences have been deposited in GenBank database under BioProject accession number PRJNA599167.

## Acknowledgements

The authors would like to thank the Smithsonian Environmental Research Center and Katrine Lohan for providing access to their facilities during sample collection. This work was supported by the National Science Foundation Biological Oceanography (awards #1820652, #1829831 and #1756314) and a Gordon and Betty Moore Foundation Investigator Award (#3790). Part of this project was conducted using computational resources at the Maryland Advanced Research Computing Center (MARCC) and the Ohio Supercomputer Center (OSC) for High Performance Computing.

## Author Contributions

EGS and SPP conceived of the work. EGS conducted all field-work, epicPCR analysis, and incubation experiments associated with this work. EGS, KAW, and SPP conducted the experimental and computational analysis for the bacterial metagenome and 16S rRNA gene libraries. EGS, FT, AAZ, and OZ conducted experimental and computational analysis for the viral metagenomic libraries. EGS, KAW, and SPP conducted the bioinformatic host prediction. EGS wrote the manuscript. EGS, AAZ, OZ, MBS, and SPP edited the manuscript.

## Additional information

*Supplementary information* is available for this paper.

## References

1 Suttle, C. A. Marine viruses—major players in the global ecosystem. Nature Reviews Microbiology 5, 801 (2007).

2 Emerson, J. B. et al. Host-linked soil viral ecology along a permafrost thaw gradient. Nature microbiology 3, 870 (2018).

3 Reyes, A., Semenkovich, N. P., Whiteson, K., Rohwer, F. & Gordon, J. I. Going viral: next-generation sequencing applied to phage populations in the human gut. Nature Reviews Microbiology 10, 607 (2012).

4 Suttle, C. A. The significance of viruses to mortality in aquatic microbial communities. Microbial ecology 28, 237–243 (1994).

5 Guidi, L. et al. Plankton networks driving carbon export in the oligotrophic ocean. Nature 532, 465–470, doi: 10.1038/nature16942 (2016).

6 Roux, S. et al. Ecogenomics and potential biogeochemical impacts of globally abundant ocean viruses. Nature 537, 689 (2016).

7 Winget, D. M. et al. Repeating patterns of virioplankton production within an estuarine ecosystem. Proceedings of the National Academy of Sciences 108, 11506–11511 (2011).

8 Chen, X. W. et al. Tide driven microbial dynamics through virus-host interactions in the estuarine ecosystem. Water Res 160, 118–129, doi: 10.1016/j.watres.2019.05.051 (2019).

9 Suttle, C. A. Marine viruses - major players in the global ecosystem. Nature Reviews Microbiology 5, 801–812, doi: 10.1038/nrmicro1750 (2007).

10 Flores, C. O., Meyer, J. R., Valverde, S., Farr, L. & Weitz, J. S. Statistical structure of host–phage interactions. Proceedings of the National Academy of Sciences 108, E288–E297 (2011).

11 Flores, C. O., Valverde, S. & Weitz, J. S. Multi-scale structure and geographic drivers of cross-infection within marine bacteria and phages. The ISME journal 7, 520–532 (2013).

12 Jover, L. F., Cortez, M. H. & Weitz, J. S. Mechanisms of multi-strain coexistence in host–phage systems with nested infection networks. Journal of theoretical biology 332, 65–77 (2013).

13 Våge, S., Storesund, J. E. & Thingstad, T. F. Adding a cost of resistance description extends the ability of virus–host model to explain observed patterns in structure and function of pelagic microbial communities. Environmental microbiology 15, 1842–1852 (2013).

14 Gregory, A. C. et al. Marine DNA Viral Macro- and Microdiversity from Pole to Pole. Cell 177, 1109-+, doi: 10.1016/j.cell.2019.03.040 (2019).

15 Tully, B. J., Graham, E. D. & Heidelberg, J. F. The reconstruction of 2,631 draft metagenome-assembled genomes from the global oceans. Scientific data 5, 170203 (2018).

16 Burstein, D. et al. Major bacterial lineages are essentially devoid of CRISPR-Cas viral defence systems. Nature communications 7, 10613 (2016).

17 Hatfull, G. F. Dark matter of the biosphere: the amazing world of bacteriophage diversity. Journal of virology 89, 8107–8110 (2015).

18 Middelboe, M., Chan, A. & Bertelsen, S. K. in Manual of aquatic viral ecology 118–133 (American Society of Limnology and Oceanography, Inc., 2010).

19 Deng, L. et al. Viral tagging reveals discrete populations in Synechococcus viral genome sequence space. Nature 513, 242-+, doi: 10.1038/nature13459 (2014).

20 Mosier-Boss, P. A. et al. Use of fluorescently labeled phage in the detection and identification of bacterial species. Appl Spectrosc 57, 1138–1144, doi:Doi 10.1366/00037020360696008 (2003).

21 Allers, E. et al. Single-cell and population level viral infection dynamics revealed by phage FISH, a method to visualize intracellular and free viruses. Environmental microbiology 15, 2306–2318 (2013).

22 Tadmor, A. D., Ottesen, E. A., Leadbetter, J. R. & Phillips, R. Probing Individual Environmental Bacteria for Viruses by Using Microfluidic Digital PCR. Science 333, 58–62, doi: 10.1126/science.1200758 (2011).

23 Spencer, S. J. et al. Massively parallel sequencing of single cells by epicPCR links functional genes with phylogenetic markers. The ISME journal 10, 427 (2016).

24 Warwick-Dugdale, J. et al. Long-read viral metagenomics captures abundant and microdiverse viral populations and their niche-defining genomic islands. PeerJ 7, e6800, doi: 10.7717/peerj.6800 (2019).

25 Brum, J. R. et al. Patterns and ecological drivers of ocean viral communities. Science 348, 1261498, doi: 10.1126/science.1261498 (2015).

26 Gregory, A. C. et al. Genomic differentiation among wild cyanophages despite widespread horizontal gene transfer. BMC Genomics 17, 930, doi: 10.1186/s12864-016-3286-x (2016).

27 Roux, S. et al. Minimum Information about an Uncultivated Virus Genome (MIUViG). Nat Biotechnol 37, 29–37, doi: 10.1038/nbt.4306 (2019).

28 Jang, H. B. et al. Taxonomic assignment of uncultivated prokaryotic virus genomes is enabled by gene-sharing networks. Nature biotechnology 37, 632–639 (2019).

29 Jasna, V., Parvathi, A. & Dash, A. Genetic and functional diversity of double-stranded DNA viruses in a tropical monsoonal estuary, India. Scientific Reports 8, 16036, doi: 10.1038/s41598-018-34332-8 (2018).

30 McDaniel, L. D., Rosario, K., Breitbart, M. & Paul, J. H. Comparative metagenomics: natural populations of induced prophages demonstrate highly unique, lower diversity viral sequences. Environmental Microbiology 16, 570–585, doi:Doi 10.1111/1462-2920.12184 (2014).

31 Allen, L. Z. et al. The Baltic Sea Virome: Diversity and Transcriptional Activity of DNA and RNA Viruses. Msystems 2, e00125–00116, doi: 10.1128/mSystems.00125-16 (2017).

32 Bench, S. R. et al. Metagenomic characterization of Chesapeake Bay virioplankton. Appl. Environ. Microbiol. 73, 7629–7641 (2007).

33 Kan, J., Evans, S. E., Chen, F. & Suzuki, M. T. Novel estuarine bacterioplankton in rRNA operon libraries from the Chesapeake Bay. Aquatic Microbial Ecology 51, 55–66 (2008).

34 Chen, F. et al. Diverse and dynamic populations of cyanobacterial podoviruses in the Chesapeake Bay unveiled through DNA polymerase gene sequences. Environmental Microbiology 11, 2884–2892, doi: 10.1111/j.1462-2920.2009.02033.x (2009).

35 Kan, J., Suzuki, M. T., Wang, K., Evans, S. E. & Chen, F. High temporal but low spatial heterogeneity of bacterioplankton in the Chesapeake Bay. Appl. Environ. Microbiol. 73, 6776–6789 (2007).

36 Nasko, D. J. et al. Family A DNA polymerase phylogeny uncovers diversity and replication gene organization in the virioplankton. Frontiers in microbiology 9, 3053 (2018).

37 Sakowski, E. G. et al. Ribonucleotide reductases reveal novel viral diversity and predict biological and ecological features of unknown marine viruses. Proceedings of the National Academy of Sciences 111, 15786–15791 (2014).

38 Adriaenssens, E. M. & Cowan, D. A. Using signature genes as tools to assess environmental viral ecology and diversity. Appl. Environ. Microbiol. 80, 4470–4480 (2014).

39 Harrison, A. O., Moore, R. M., Polson, S. W. & Wommack, K. E. Reannotation of the ribonucleotide reductase in a cyanophage reveals life history strategies within the virioplankton. Frontiers in microbiology 10, 134 (2019).

40 Martinez-Hernandez, F. et al. Single-virus genomics reveals hidden cosmopolitan and abundant viruses. Nature communications 8, 15892 (2017).

41 Galiez, C., Siebert, M., Enault, F., Vincent, J. & Soding, J. WIsH: who is the host? Predicting prokaryotic hosts from metagenomic phage contigs. Bioinformatics 33, 3113–3114, doi: 10.1093/bioinformatics/btx383 (2017).

42 Galiez, C., Siebert, M., Enault, F., Vincent, J. & Söding, J. WIsH: who is the host? Predicting prokaryotic hosts from metagenomic phage contigs. Bioinformatics 33, 3113–3114 (2017).

43 Kavagutti, V. S., Andrei, A. S., Mehrshad, M., Salcher, M. M. & Ghai, R. Phage-centric ecological interactions in aquatic ecosystems revealed through ultra-deep metagenomics. Microbiome 7, 135, doi: 10.1186/s40168-019-0752-0 (2019).

44 Holmfeldt, K., Middelboe, M., Nybroe, O. & Riemann, L. Large variabilities in host strain susceptibility and phage host range govern interactions between lytic marine phages and their Flavobacterium hosts. Appl. Environ. Microbiol. 73, 6730–6739 (2007).

45 Tzortziou, M. et al. Tidal marshes as a source of optically and chemically distinctive colored dissolved organic matter in the Chesapeake Bay. Limnology and Oceanography 53, 148–159, doi:DOI 10.4319/lo.2008.53.1.0148 (2008).

46 Chai, T. J. Characteristics of Escherichia-Coli Grown in Bay Water as Compared with Rich Medium. Applied and Environmental Microbiology 45, 1316–1323, doi:Doi 10.1128/Aem.45.4.1316-1323.1983 (1983).

47 Martiny, J. B., Riemann, L., Marston, M. F. & Middelboe, M. Antagonistic coevolution of marine planktonic viruses and their hosts. (2014).

48 Sieradzki, E. T., Ignacio-Espinoza, J. C., Needham, D. M., Fichot, E. B. & Fuhrman, J. A. Dynamic marine viral infections and major contribution to photosynthetic processes shown by spatiotemporal picoplankton metatranscriptomes. Nat Commun 10, 1169, doi: 10.1038/s41467-019-09106-z (2019).

49 Moniruzzaman, M. et al. Virus-host relationships of marine single-celled eukaryotes resolved from metatranscriptomics. Nat Commun 8, 16054, doi: 10.1038/ncomms16054 (2017).

50 Duffy, S., Turner, P. E. & Burch, C. L. Pleiotropic costs of niche expansion in the RNA bacteriophage F6. Genetics 172, 751–757 (2006).

51 Sunagawa, S. et al. Structure and function of the global ocean microbiome. Science 348, 1261359, doi: 10.1126/science.1261359 (2015).

52 Roux, S. et al. Ecogenomics and potential biogeochemical impacts of globally abundant ocean viruses. Nature 537, 689-+, doi: 10.1038/nature19366 (2016).

53 Deng, L. et al. Contrasting life strategies of viruses that infect photo-and heterotrophic bacteria, as revealed by viral tagging. MBio 3, e00373–00312 (2012).

54 Labonté, J. M. et al. Single-cell genomics-based analysis of virus–host interactions in marine surface bacterioplankton. The ISME journal 9, 2386–2399 (2015).

55 Adriaenssens, E. M. & Cowan, D. A. Using Signature Genes as Tools To Assess Environmental Viral Ecology and Diversity. Applied and Environmental Microbiology 80, 4470–4480, doi: 10.1128/Aem.00878-14 (2014).

56 Martinez-Hernandez, F. et al. Single-virus genomics reveals hidden cosmopolitan and abundant viruses. Nat Commun 8, 15892, doi: 10.1038/ncomms15892 (2017).

57 Martinez-Hernandez, F. et al. Droplet Digital PCR for Estimating Absolute Abundances of Widespread Pelagibacter Viruses. Front Microbiol 10, 1226, doi: 10.3389/fmicb.2019.01226 (2019).

58 Vik, D. R. et al. Putative archaeal viruses from the mesopelagic ocean. PeerJ 5, e3428, doi: 10.7717/peerj.3428 (2017).

59 Jover, L. F., Romberg, J. & Weitz, J. S. Inferring phage–bacteria infection networks from time-series data. Royal Society Open Science 3, 160654 (2016).

60 Labonte, J. M. et al. Single-cell genomics-based analysis of virus-host interactions in marine surface bacterioplankton. Isme J 9, 2386–2399, doi: 10.1038/ismej.2015.48 (2015).

61 Brankatschk, R., Bodenhausen, N., Zeyer, J. & Burgmann, H. Simple Absolute Quantification Method Correcting for Quantitative PCR Efficiency Variations for Microbial Community Samples. Applied and Environmental Microbiology 78, 4481–4489, doi: 10.1128/Aem.07878-11 (2012).

62 Baran, N., Goldin, S., Maidanik, I. & Lindell, D. Quantification of diverse virus populations in the environment using the polony method. Nature Microbiology 3, doi: 10.1038/s41564-017-0045-y (2018).

63 Russell, D. A. & Hatfull, G. F. PhagesDB: the actinobacteriophage database. Bioinformatics 33, 784–786, doi: 10.1093/bioinformatics/btw711 (2017).

64 Jensen, E. C. et al. Prevalence of broad-host-range lytic bacteriophages of Sphaerotilus natans, Escherichia coli, and Pseudomonas aeruginosa. Applied and Environmental Microbiology 64, 575–580 (1998).

65 Peters, D. L., Lynch, K. H., Stothard, P. & Dennis, J. J. The isolation and characterization of two Stenotrophomonas maltophilia bacteriophages capable of cross-taxonomic order infectivity. BMC Genomics 16, 664, doi: 10.1186/s12864-015-1848-y (2015).

66 Paez-Espino, D. et al. Uncovering Earth’s virome. Nature 536, 425-+, doi: 10.1038/nature19094 (2016).

67 Chiura, H. X. Generalized gene transfer by virus-like particles from marine bacteria. Aquatic Microbial Ecology 13, 75–83 (1997).

68 Hanski, I., Hansson, L. & Henttonen, H. SPECIALIST PREDATORS, GENERALIST PREDATORS, AND THE MICROTINE RODENT CYCLE. Journal of Animal Ecology 60, 353–367, doi: 10.2307/5465 (1991).

69 Johnke, J. et al. A Generalist Protist Predator Enables Coexistence in Multitrophic Predator-Prey Systems Containing a Phage and the Bacterial Predator Bdellovibrio. Front Ecol Evol 5 (2017).

70 John, S. G. et al. A simple and efficient method for concentration of ocean viruses by chemical flocculation. Environmental microbiology reports 3, 195–202 (2011).

71 Bolger, A. M., Lohse, M. & Usadel, B. Trimmomatic: a flexible trimmer for Illumina sequence data. Bioinformatics 30, 2114–2120 (2014).

72 Bankevich, A. et al. SPAdes: A New Genome Assembly Algorithm and Its Applications to Single-Cell Sequencing. Journal of Computational Biology 19, 455–477, doi: 10.1089/cmb.2012.0021 (2012).

73 Uritskiy, G. V., DiRuggiero, J. & Taylor, J. MetaWRAP—a flexible pipeline for genome-resolved metagenomic data analysis. Microbiome 6, 158 (2018).

74 Kang, D. D., Froula, J., Egan, R. & Wang, Z. MetaBAT, an efficient tool for accurately reconstructing single genomes from complex microbial communities. PeerJ 3, e1165 (2015).

75 Wu, Y.-W., Simmons, B. A. & Singer, S. W. MaxBin 2.0: an automated binning algorithm to recover genomes from multiple metagenomic datasets. Bioinformatics 32, 605–607 (2015).

76 Song, W.-Z. & Thomas, T. Binning_refiner: improving genome bins through the combination of different binning programs. Bioinformatics 33, 1873–1875 (2017).

77 Parks, D. H., Imelfort, M., Skennerton, C. T., Hugenholtz, P. & Tyson, G. W. CheckM: assessing the quality of microbial genomes recovered from isolates, single cells, and metagenomes. Genome research 25, 1043–1055 (2015).

78 Menzel, P., Ng, K. L. & Krogh, A. Fast and sensitive taxonomic classification for metagenomics with Kaiju. Nat Commun 7, 11257, doi: 10.1038/ncomms11257 (2016).

79 Schütze, T. et al. A streamlined protocol for emulsion polymerase chain reaction and subsequent purification. Analytical biochemistry 410, 155–157 (2011).

80 Bolyen, E. et al. QIIME 2: Reproducible, interactive, scalable, and extensible microbiome data science. Report No. 2167-9843, (PeerJ Preprints, 2018).

81 DeSantis, T. Z. et al. Greengenes, a chimera-checked 16S rRNA gene database and workbench compatible with ARB. Appl. Environ. Microbiol. 72, 5069–5072 (2006).

82 Hurwitz, B. L., Deng, L., Poulos, B. T. & Sullivan, M. B. Evaluation of methods to concentrate and purify ocean virus communities through comparative, replicated metagenomics. Environmental Microbiology 15, 1428–1440 (2013).

83 Bushnell, B. BBMap: A Fast, Accurate, Splice-Aware Aligner, <https://www.osti.gov/servlets/purl/1241166> (2014).

84 De Coster, W., D’Hert, S., Schultz, D. T., Cruts, M. & Van Broeckhoven, C. NanoPack: visualizing and processing long-read sequencing data. Bioinformatics 34, 2666–2669, doi: 10.1093/bioinformatics/bty149 (2018).

85 Kolmogorov, M., Yuan, J., Lin, Y. & Pevzner, P. A. Assembly of long, error-prone reads using repeat graphs. Nat Biotechnol 37, 540-+, doi: 10.1038/s41587-019-0072-8 (2019).

86 Walker, B. J. et al. Pilon: An Integrated Tool for Comprehensive Microbial Variant Detection and Genome Assembly Improvement. Plos One 9, e112963, doi: 10.1371/journal.pone.0112963 (2014).

87 Li, H. & Durbin, R. Fast and accurate short read alignment with Burrows-Wheeler transform. Bioinformatics 25, 1754–1760, doi: 10.1093/bioinformatics/btp324 (2009).

88 Roux, S., Enault, F., Hurwitz, B. L. & Sullivan, M. B. VirSorter: mining viral signal from microbial genomic data. PeerJ 3, e985, doi: 10.7717/peerj.985 (2015).

89 Ren, J. et al. Identifying viruses from metagenomic data using deep learning. Quantitative Biology, doi: 10.1007/s40484-019-0187-4 (2020).

90 Bolduc, B., Youens-Clark, K., Roux, S., Hurwitz, B. L. & Sullivan, M. B. iVirus: facilitating new insights in viral ecology with software and community data sets imbedded in a cyberinfrastructure. Isme J 11, 7–14, doi: 10.1038/ismej.2016.89 (2017).

91 Langmead, B. & Salzberg, S. L. Fast gapped-read alignment with Bowtie 2. Nat Methods 9, 357–U354, doi: 10.1038/Nmeth.1923 (2012).

92 Jang, H. B. et al. Taxonomic assignment of uncultivated prokaryotic virus genomes is enabled by gene-sharing networks. Nat Biotechnol 37, 632-+, doi: 10.1038/s41587-019-0100-8 (2019).

93 Hyatt, D. et al. Prodigal: prokaryotic gene recognition and translation initiation site identification. BMC Bioinformatics 11, 119, doi: 10.1186/1471-2105-11-119 (2010).

94 Smoot, M. E., Ono, K., Ruscheinski, J., Wang, P. L. & Ideker, T. Cytoscape 2.8: new features for data integration and network visualization. Bioinformatics 27, 431–432, doi: 10.1093/bioinformatics/btq675 (2011).

95 Caporaso, J. G. et al. QIIME allows analysis of high-throughput community sequencing data. Nature methods 7, 335 (2010).

96 Warnes, G. R. et al. gplots: Various R programming tools for plotting data. (2015).

97 Langmead, B. & Salzberg, S. L. Fast gapped-read alignment with Bowtie 2. Nature methods 9, 357 (2012).

98 Zhu, W., Lomsadze, A. & Borodovsky, M. Ab initio gene identification in metagenomic sequences. Nucleic acids research 38, e132–e132 (2010).

99 Suzek, B. E., Huang, H., McGarvey, P., Mazumder, R. & Wu, C. H. UniRef: comprehensive and non-redundant UniProt reference clusters. Bioinformatics 23, 1282–1288 (2007).

100 Katoh, K. & Standley, D. M. MAFFT multiple sequence alignment software version 7: improvements in performance and usability. Molecular biology and evolution 30, 772–780 (2013).

101 Kearse, M. et al. Geneious Basic: an integrated and extendable desktop software platform for the organization and analysis of sequence data. Bioinformatics 28, 1647–1649 (2012).

102 Martin, M. Cutadapt removes adapter sequences from high-throughput sequencing reads. EMBnet. journal 17, 10–12 (2011).

103 Fu, L., Niu, B., Zhu, Z., Wu, S. & Li, W. CD-HIT: accelerated for clustering the next-generation sequencing data. Bioinformatics 28, 3150–3152 (2012).

104 Dereeper, A. et al. Phylogeny. fr: robust phylogenetic analysis for the non-specialist. Nucleic acids research 36, W465–W469 (2008).

