## Supplemental Information for "Capturing *in situ* Virus-Host Range and Interaction Dynamics through Gene Fusion with epicPCR"

*Sakowski et al.*

### SUPPLEMENTAL TABLES

**Table S1.** Top BLASTn hits for the Chesapeake Bay 'Cyano SP'-like phage RNR gene sequences.

| Phage Cluster ID | Phage Clade | Top Hit in NCBI nr database (% ID) | Top Reference Hit (% ID) |
|---|---|---|---|
| **25859** | I | Synechococcus phage S-CBP3 (78%) | Synechococcus phage S-CBP3 (78%) |
| **26558** | I | Synechococcus phage S-CBP3 (78%) | Synechococcus phage S-CBP3 (78%) |
| **26599** | I | Synechococcus phage S-CBP3 (78%) | Synechococcus phage S-CBP3 (78%) |
| **26609** | I | Synechococcus phage S-CBP3 (78%) | Synechococcus phage S-CBP3 (78%) |
| **26972** | I | Synechococcus phage S-CBP3 (78%) | Synechococcus phage S-CBP3 (78%) |
| **3204** | I | Synechococcus phage S-CBP3 (78%) | Synechococcus phage S-CBP3 (78%) |
| **11846** | II | Uncultured virus clone CFB_Contig_5 (89%) | Synechococcus phage S-CBP4 (71%) |
| **12891** | II | Uncultured virus clone CFB_Contig_5 (89%) | Synechococcus phage S-CBP4 (71%) |
| **5666** | II | Uncultured virus clone CFB_Contig_5 (89%) | Synechococcus phage S-RIP1 (71%) |
| **6908** | II | Uncultured virus clone CFB_Contig_5 (90%) | Synechococcus phage S-CBP4 (71%) |
| **6971** | II | Uncultured virus clone CFB_Contig_5 (89%) | Synechococcus phage S-CBP4 (71%) |
| **7860** | II | Uncultured virus clone CFB_Contig_5 (89%) | Synechococcus phage S-CBP4 (72%) |
| **8094** | II | Uncultured virus clone CFB_Contig_5 (90%) | Synechococcus phage S-CBP4 (71%) |
| **9966** | II | Uncultured virus clone CFB_Contig_5 (89%) | Synechococcus phage S-CBP4 (72%) |
| **11061** | III | Uncultured virus clone CFB_Contig_5 (85%) | Synechococcus phage S-CBP1 (72%) |
| **2950** | III | Uncultured virus clone CFB_Contig_5 (85%) | Synechococcus phage S-CBP1 (72%) |
| **4035** | III | Uncultured virus clone CFB_Contig_5 (85%) | Synechococcus phage S-CBP1 (72%) |
| **4275** | III | Uncultured virus clone CFB_Contig_5 (85%) | Synechococcus phage S-CBP1 (72%) |

| | | | |
|---|---|---|---|
| **4655** | III | Uncultured virus clone CFB_Contig_5 (85%) | Synechococcus phage S-CBP1 (72%) |
| **4870** | III | Uncultured virus clone CFB_Contig_5 (85%) | Synechococcus phage S-CBP1 (72%) |
| **5531** | III | Uncultured virus clone CFB_Contig_5 (85%) | Synechococcus phage S-RIP1 (72%) |
| **6666** | III | Uncultured virus clone CFB_Contig_5 (85%) | Synechococcus phage S-CBP1 (72%) |
| **7153** | III | Uncultured virus clone CFB_Contig_5 (85%) | Synechococcus phage S-CBP1 (72%) |
| **7184** | III | Uncultured virus clone CFB_Contig_5 (85%) | Synechococcus phage S-CBP1 (72%) |
| **7194** | III | Uncultured virus clone CFB_Contig_5 (86%) | Synechococcus phage S-CBP1 (72%) |
| **7233** | III | Uncultured virus clone CFB_Contig_5 (85%) | Synechococcus phage S-CBP1 (72%) |
| **7304** | III | Uncultured virus clone CFB_Contig_5 (85%) | Synechococcus phage S-CBP1 (72%) |
| **7330** | III | Uncultured virus clone CFB_Contig_5 (86%) | Synechococcus phage S-CBP1 (72%) |
| **7526** | III | Uncultured virus clone CFB_Contig_5 (85%) | Synechococcus phage S-CBP1 (72%) |
| **7659** | III | Uncultured virus clone CFB_Contig_5 (85%) | Synechococcus phage S-CBP1 (72%) |
| **7704** | III | Uncultured virus clone CFB_Contig_5 (85%) | Synechococcus phage S-CBP1 (72%) |
| **8414** | III | Uncultured virus clone CFB_Contig_5 (85%) | Synechococcus phage S-RIP1 (72%) |
| **9769** | III | Uncultured virus clone CFB_Contig_5 (86%) | Synechococcus phage S-CBP1 (72%) |
| **26502** | N/A | Synechococcus phage S-CBP4 (77%) | Synechococcus phage S-CBP4 (77%) |
| **26534** | N/A | Synechococcus phage S-CBP4 (77%) | Synechococcus phage S-CBP4 (77%) |
| **26870** | N/A | Synechococcus phage S-CBP4 (78%) | Synechococcus phage S-CBP4 (78%) |
| **8027** | N/A | Uncultured virus clone CFB_Contig_5 (86%) | Synechococcus phage S-CBP1 (71%) |
| **8065** | N/A | Uncultured virus clone CFB_Contig_5 (94%) | Synechococcus phage S-CBP4 (71%) |
| **8091** | N/A | Uncultured virus clone CFB_Contig_5 (95%) | Synechococcus phage S-CBP3 (72%) |
| **8372** | N/A | Uncultured virus clone CFB_Contig_5 (94%) | Synechococcus phage S-CBP4 (71%) |

**Table S2.** Spearman's Rank correlations between the Chesapeake Bay physicochemical properties and 'Cyano-SP'-like phage abundances and phage-host interactions.

| | Abundance | Clade I Interactions | Clade II Interactions | Clade III Interactions |
|---|---|---|---|---|
| **Temperature (°C)** | $p < 0.05$ (+) | NS | $p < 0.001$ (+) | $p = 0.02$ (+) |
| **Salinity (psu)** | $p < 0.01$ (+) | NS | NS | $p = 0.02$ (+) |
| **pH** | NS | NS | $p = 0.02$ (-) | NS |
| **Chlorophyll (RFU)** | NS | $p = 0.005$ (+) | NS | NS |
| **Dissolved Oxygen (mg/L)** | NS | NS | $p < 0.001$ (-) | NS |
| **Turbidity (FNU)** | NS | NS | $p = 0.01$ (+) | NS |
| **FDOM (RFU)** | NS | NS | NS | $p = 0.003$ (-) |
| **Water Level (mm)** | NS | NS | NS | NS |

**Table S3.** Chesapeake Bay surface water physicochemical measurements at the Smithsonian Environmental Research Center.

| Sample Date | Water Temperature (°C) | Water pH | Water Chlorophyll (RFU) | Water Total Algae (RFU) | Fluorescent Dissolved Organic Matter (RFU) | Water Dissolved Oxygen (mg $L^{-1}$) | Probe Depth (m) | Water Turbidity (FNU) |
|---|---|---|---|---|---|---|---|---|
| **5/24/18** | 23.72 | 8.4 | 9.9 | 20.6 | 13.2 | 9.19 | 1.09 | 7.2 |
| **5/31/18** | 25.37 | 8.1 | 26.1 | 45.4 | 12.7 | 8.28 | 1.01 | 12.16 |
| **6/8/18** | 24.4 | 8.6 | 22.1 | 41.8 | 11.8 | 10.34 | 1.08 | 13.12 |
| **6/14/18** | 24.82 | 7.7 | 7.4 | 12.3 | 10.2 | 6.34 | 1 | 14.34 |
| **6/22/18** | 26.51 | 7.4 | 8 | 23.6 | 9 | 5.33 | 0.98 | 11.66 |
| **6/28/18** | 25.86 | 7.6 | 7.9 | 11.3 | 8.9 | 6.73 | 0.66 | |
| **7/5/18** | 31.64 | 7.7 | 7.7 | 11.8 | 8.1 | 6.52 | 0.82 | 9.76 |
| **7/12/18** | 29.5 | 8.1 | 12.9 | 22.5 | 8.9 | 8.32 | 0.78 | 13.87 |
| **7/19/18** | 28.69 | 7.7 | 11.1 | 12.3 | 8.9 | 5.91 | 0.76 | 17.66 |
| **7/26/18** | 26.3 | 7.3 | 6.4 | 10.5 | 15.7 | 3.99 | 0.7 | 15.61 |
| **8/2/18** | 28.34 | 8.6 | 16.2 | 31.4 | 14 | 9.43 | 1.04 | 18.57 |
| **8/9/18** | 30 | 7.9 | 5.4 | 14.3 | 12.4 | 4.44 | 0.94 | 17.07 |
| **8/23/18** | 25.83 | 7.9 | 13.7 | 31.2 | 13 | 5.92 | 1.03 | 27.71 |
| **12/6/18** | 6.14 | 7.8 | 1.9 | 4 | 12 | 10.63 | 1.12 | 6.95 |
| **12/14/18** | 4.94 | 7.97 | 4.3 | 8 | 11.2 | 12.14 | 1.16 | 5.72 |
| **12/20/18** | 5.5 | 8.18 | 6.3 | 10.2 | 13.1 | 13 | 1.03 | 5.67 |
| **12/27/18** | 7.36 | 8.51 | 14.5 | 29.7 | 10.7 | 13.05 | 1.21 | 9.14 |

**Table S4**. Primer sequences used in this study.

| Primer Name | Gene Target | Sequence (5’ - 3’) | Citations |
|---|---|---|---|
| Cyano_SP_F primer | Ribonucleotide reductase | GGWGAYATYTGGCTMAACAA | This study |
| Cyano_SP_R_519R | Ribonucleotide reductase | R1: GWATTACCGCGGCKGCTGTTGAAGCTRTADCCRTGMAGA<br>R2: GWATTACCGCGGCKGCTGTTRAATGARTATCCRTGRAGA | This study |
| Cyano_SP_Nested_F A | Ribonucleotide reductase | F1: AAYGTCTGCCTTGARGTTT<br>F2: AAYGTWTGCCTGGAGGTGT | This study |
| Cyano_SP_Nested_F B | Ribonucleotide reductase | F1: ARGTWTACYTSCCSTCACGM<br>F2: ARGTKTAYYTGCCWWCACGR | This study |
| 27F | 16S rRNA | AGAGTTTGATCMTGGCTCAG | Lane, 1991 |
| PE16S_V4_U515_F | 16S rRNA | ACACGACGCTCTTCCGATCTYRYRGTGCCAGCMGCCGCGGTAA | Preheim et al. 2013 |
| PE_16S_V4_E786_R | 16S rRNA | CGGCATTCCTGCTGAACCGCTCTTCCGATCTGGACTACHVGGGTWT CTAAT | Preheim et al. 2013 |
| S-*-Univ-1100-a-A-15 | 16S rRNA | GGGTYKCGCTCGTTR | Klindworth et al. 2013 |
| 1492R | 16S rRNA | TACGGYTACCTTGTTACGACTT | Lane, 1991 |
| U519F-block10 | N/A | TTTTTTTTTTCAGCMGCCGCGGTAATWC/3SpC3/ | Spencer et al. 2016 |
| U519R-block10 | N/A | TTTTTTTTTTGWATTACCGCGGCKGCTG/3SpC3/ | Spencer et al. 2016 |
| PE-III-PCR-F | Illumina Forward Adapter | AATGATACGGCGACCACCGAGATCTACACTCTTTCCCTACACGACG CTCTTCCGATCT | |
| PE-IV-PCR-XXX | 16S Barcode with Illumina Reverse Adapter | CAAGCAGAAGACGGCATACGAGAT-Barcode-CGGTCTCGGCATTCCTGCTGAACCGCTCTTCCGATCT | Preheim et al. 2016 |
| Barcode_Rev | Illumina Reverse Adapter | CAAGCAGAAGACGGCATACGAGAT | |

### SUPPLEMENTAL FIGURES

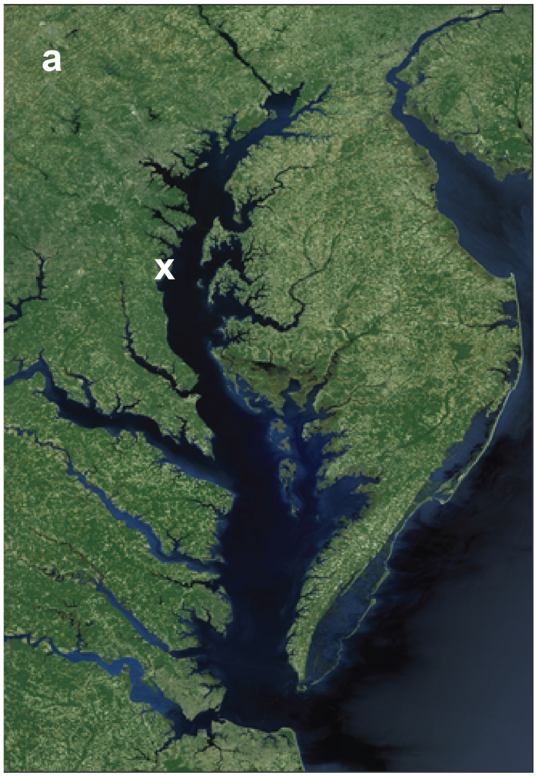


b

| Date | epicPCR | 16S rRNA gene | Virome | Metagenome | Incubation | SAG |
|---|---|---|---|---|---|---|
| 5/17/17 | | ■ | ■ | ■ | | |
| 5/18/17 | | | | ■ | | ■ |
| 5/19/17 | | | | ■ | | |
| 5/24/18 | ■ | ■ | | | | |
| 5/31/18 | ■ | ■ | ■ | ■ | | |
| 6/7/18 | ■ | ■ | | | | |
| 6/14/18 | ■ | ■ | | | | |
| 6/21/18 | ■ | ■ | | | | |
| 6/28/18 | ■ | ■ | | ■ | | |
| 7/5/18 | ■ | ■ | | | | |
| 7/12/18 | ■ | ■ | | | | |
| 7/19/18 | ■ | ■ | | | | |
| 7/26/18 | ■ | ■ | | | | |
| 8/2/18 | ■ | ■ | ■ | ■ | | |
| 8/9/18 | ■ | ■ | | | | |
| 8/23/18 | ■ | ■ | | | | |
| 12/6/18 | ■ | ■ | | ■ | | |
| 12/14/18 | ■ | ■ | ■ | | | |
| 12/20/18 | ■ | ■ | ■ | | | |
| 12/27/18 | ■ | ■ | | | | |
| 7/25/19 | | | | | ■ | |

**Figure S1.** Overview of sampling site and sample collection and analysis. a.) Map of the Chesapeake Bay with sampling site at the Smithsonian Environmental Research Center (SERC) in Edgewater, MD marked with an X. Samples were taken off the SERC pier near the mouth of the Rhode River. b.) Sampling time-line and corresponding experimental method(s) applied to samples collected from each time period. SAG: single-cell genomics.

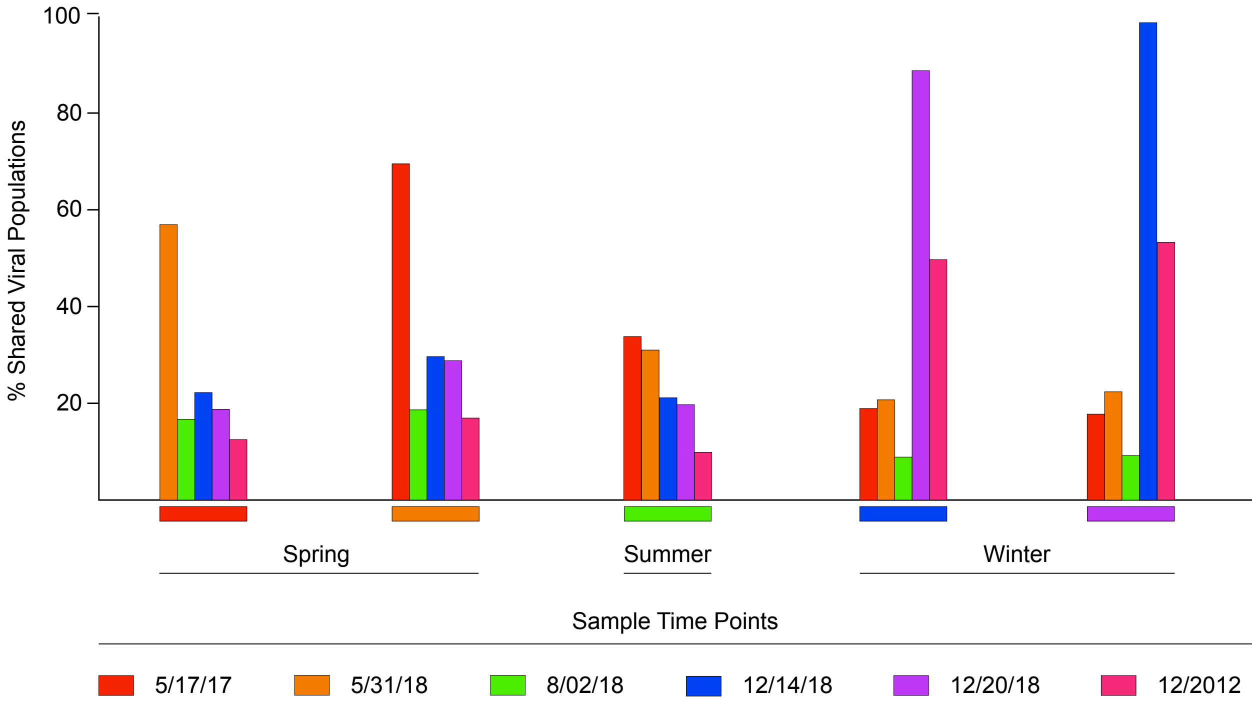


**Figure S2**. Shared viral populations between spring, summer, and winter Chesapeake Bay viral communities. Viral populations were defined as contigs >5kb with < 95% average nucleotide identity across 80% of the contig. Vertical and horizontal (below each bar cluster) bar color represent the date of sampling for each group, according to the figure legend.

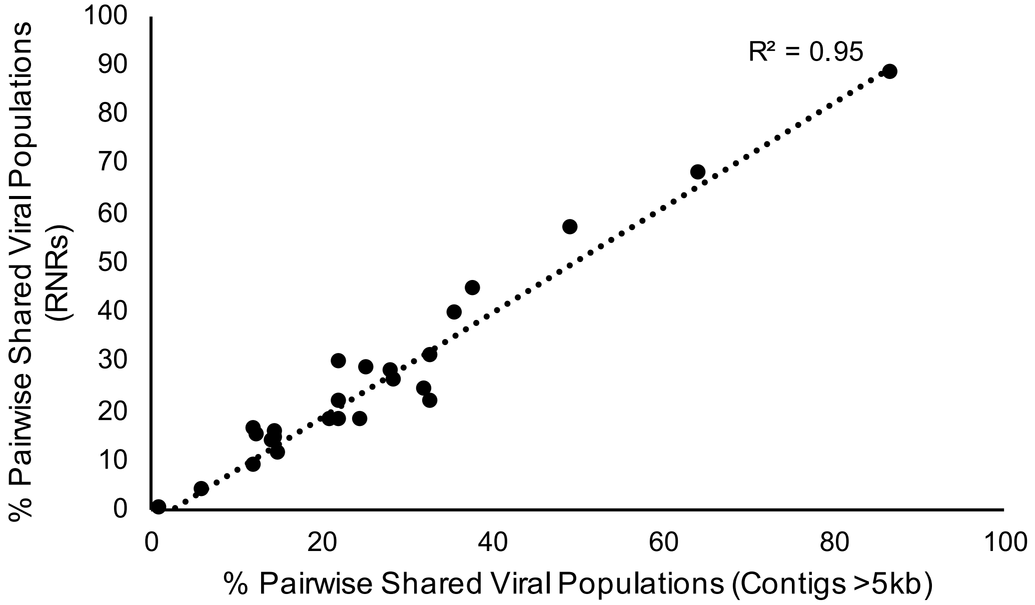


**Figure S3.** Correlation of pairwise shared populations determined by comparing viral contigs and alpha subunit RNR genes. RNR genes alone captured the seasonal diversity observed from viral contigs, making it a good marker gene for viral population diversity.

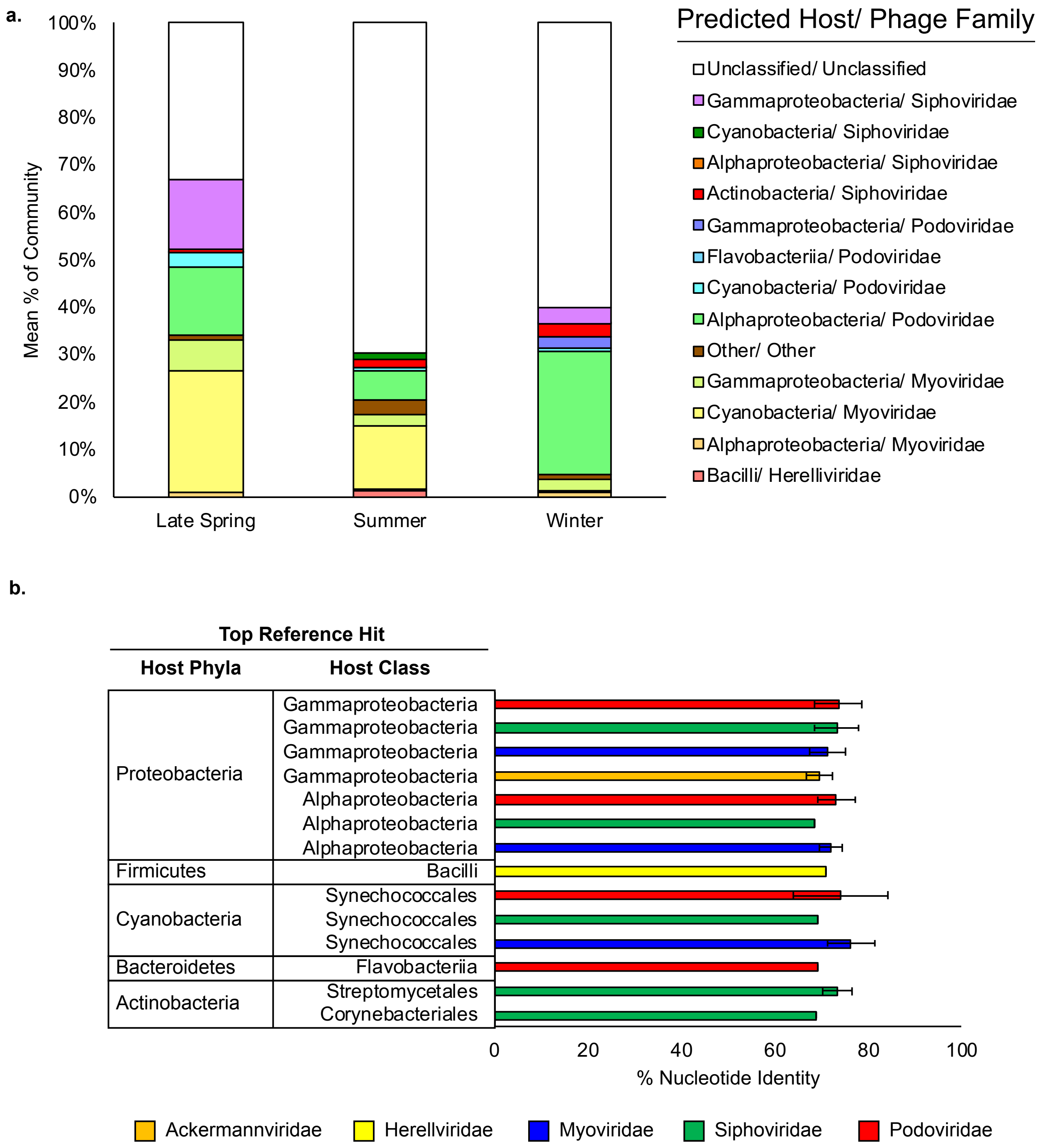


**Figure S4**. Host predictions for Chesapeake Bay viral populations from spring (May), summer (August), and winter (December) virome libraries. A) Predicted host and phage family based on RNR homology. RNR homology was queried by BLASTn against the NCBI nr database. Only top hits with e values < 1E-10 were classified. B) Nucleotide identity between Chesapeake Bay RNR sequences and reference sequence top hits. Error bars are standard deviation.

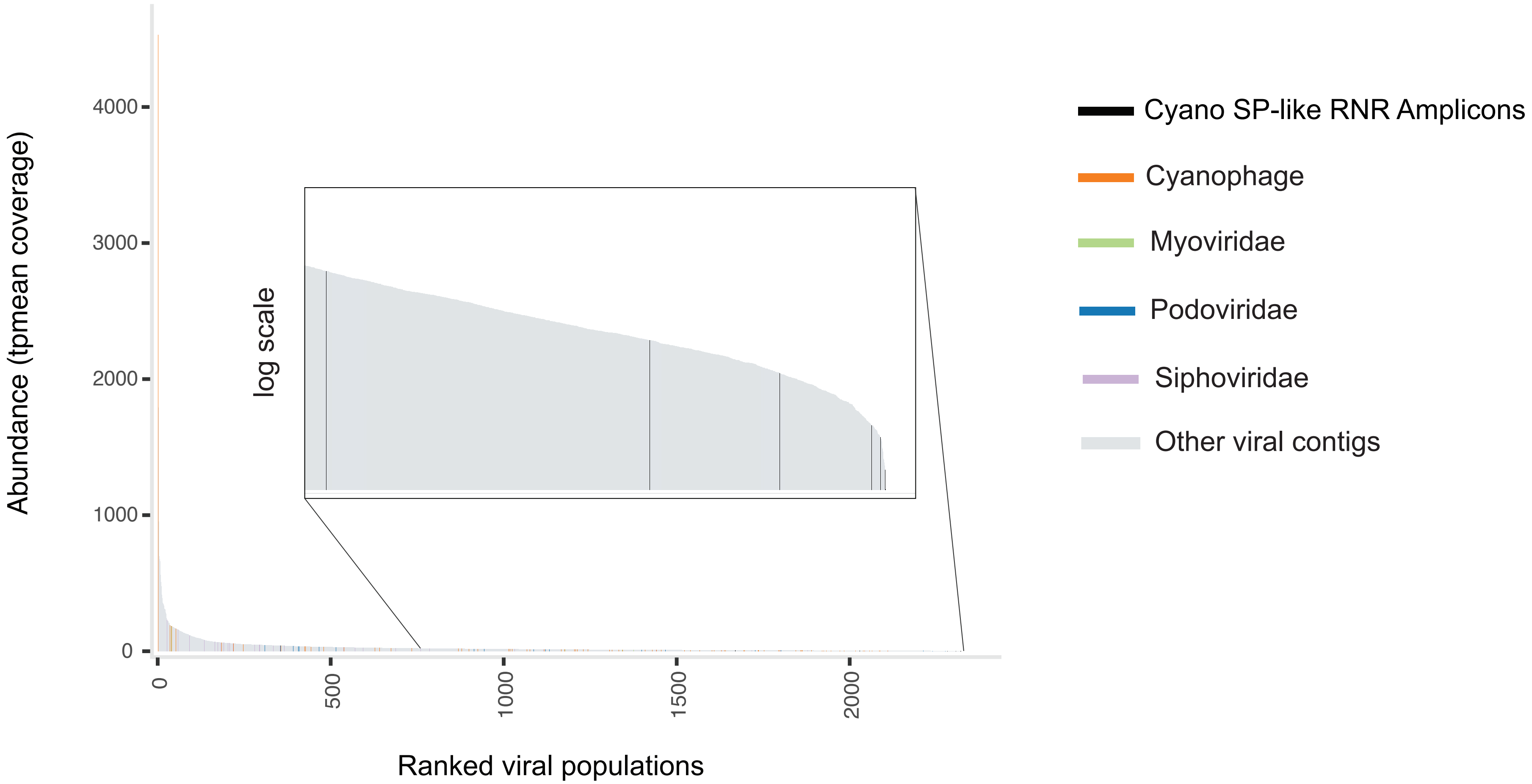


**Figure S5**. Rank abundance curve of viral populations and RNR sequences in an example viral community. All of the RNR sequences were ranked in the rare tails of the two viral communities and are shown in the blown-out, log-transformed insets. Only the two paired log-read and short-read metagenomes are shown here with similar results being observed for the rest of the short-read-only viromes.

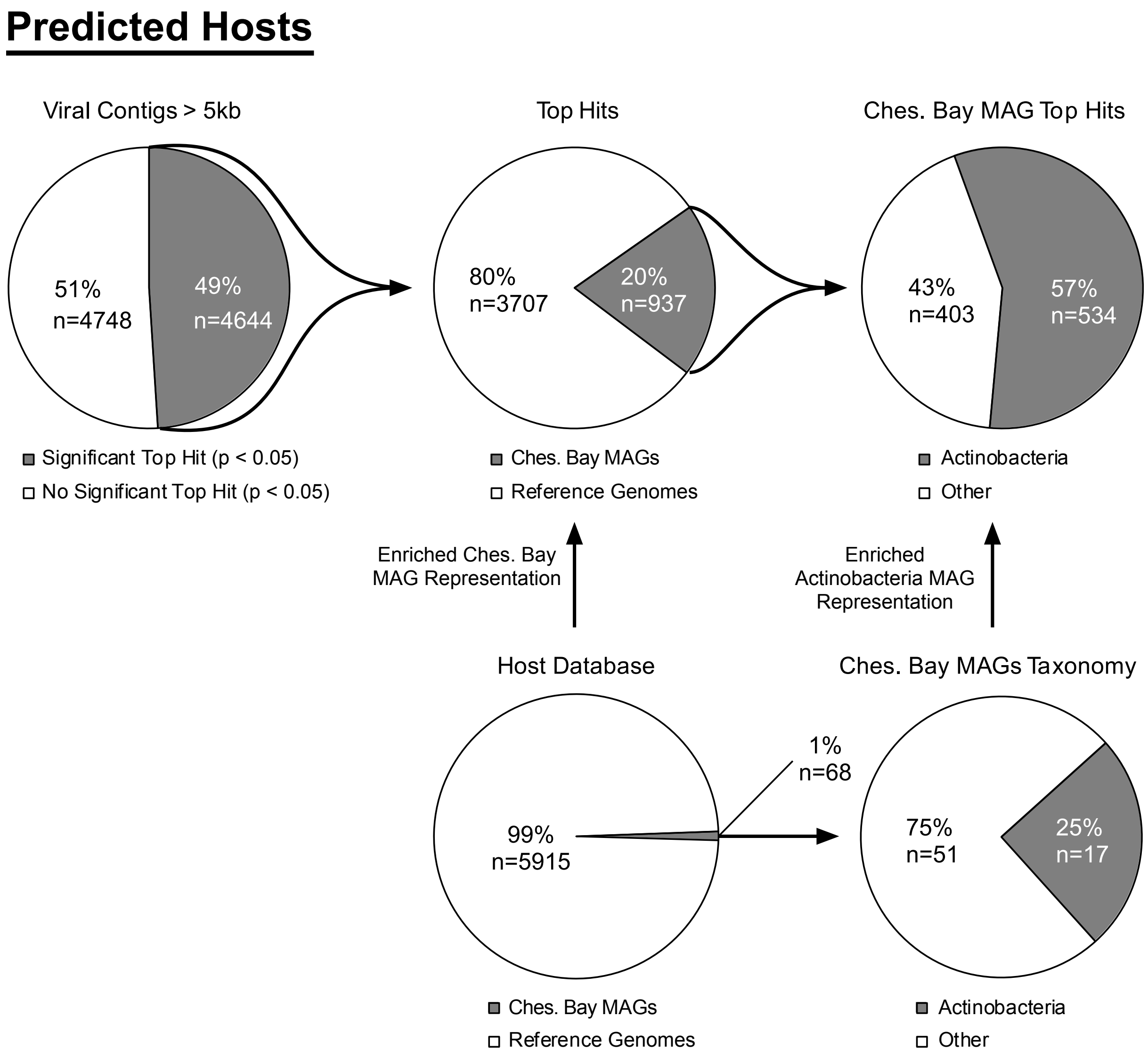


**Figure S6.** Kmer-based host predictions for Chesapeake Bay viral contigs assembled from shotgun metagenomics sequence data. Half of all assembled viral contigs > 10kb had a significant top hit to a putative host in the host database (see Materials and Methods; upper panel left). Actinobacteria were overrepresented as putative hosts for Chesapeake Bay viral contigs relative to all Chesapeake Bay metagenome-assembled genomes (MAGs; upper panel right). If the composition of predicted top hosts and the abundance of those in database were very similar, it would suggest that the probability of being a predicted host would scale with the abundance in the database, potentially creating false-positive associations. However, the enrichment of MAGs in the top host predictions (upper panel middle) compared to the number of MAGs in the database (lower panel left) and the enrichment of Actinobacteria within MAGs predicted as hosts (upper panel right) compared to the composition of the MAG dataset (lower panel right) is consistent with substantial viral pressure on Actinobacteria populations in this environment.

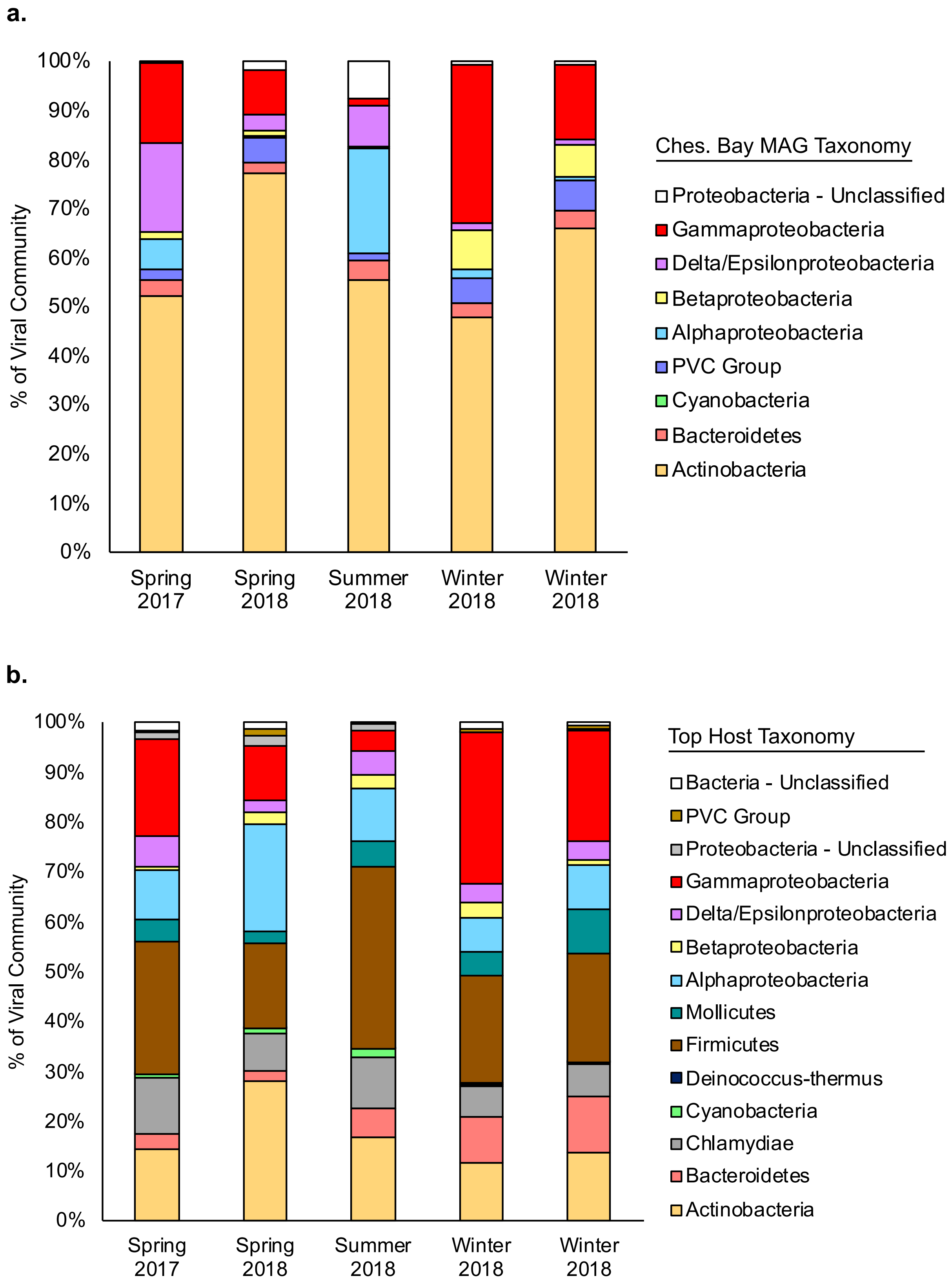
a.
100%
90%
80%
70%
60%
50%
40%
30%
20%
10%
0%
% of Viral Community
Spring 2017
Spring 2018
Summer 2018
Winter 2018
Winter 2018
Ches. Bay MAG Taxonomy
Proteobacteria - Unclassified
Gammaproteobacteria
Delta/Epsilonproteobacteria
Betaproteobacteria
Alphaproteobacteria
PVC Group
Cyanobacteria
Bacteroidetes
Actinobacteria
b.
100%
90%
80%
70%
60%
50%
40%
30%
20%
10%
0%
% of Viral Community
Spring 2017
Spring 2018
Summer 2018
Winter 2018
Winter 2018
Top Host Taxonomy
Bacteria - Unclassified
PVC Group
Proteobacteria - Unclassified
Gammaproteobacteria
Delta/Epsilonproteobacteria
Betaproteobacteria
Alphaproteobacteria
Mollicutes
Firmicutes
Deinococcus-thermus
Cyanobacteria
Chlamydiae
Bacteroidetes
Actinobacteria

**Figure S7.** Viral community composition of contigs based on top predicted host by *in silico* host prediction. A) Viral community composition of contigs with a Chesapeake Bay MAG as a top predicted host. Contigs with a Chesapeake Bay MAG as a top predicted host represented 20% of all viral contigs with a significant ($p < 0.05$) host prediction ($n = 937$). B) Viral community composition of all contigs with a significant ($p < 0.05$) host prediction ($n = 4,644$).

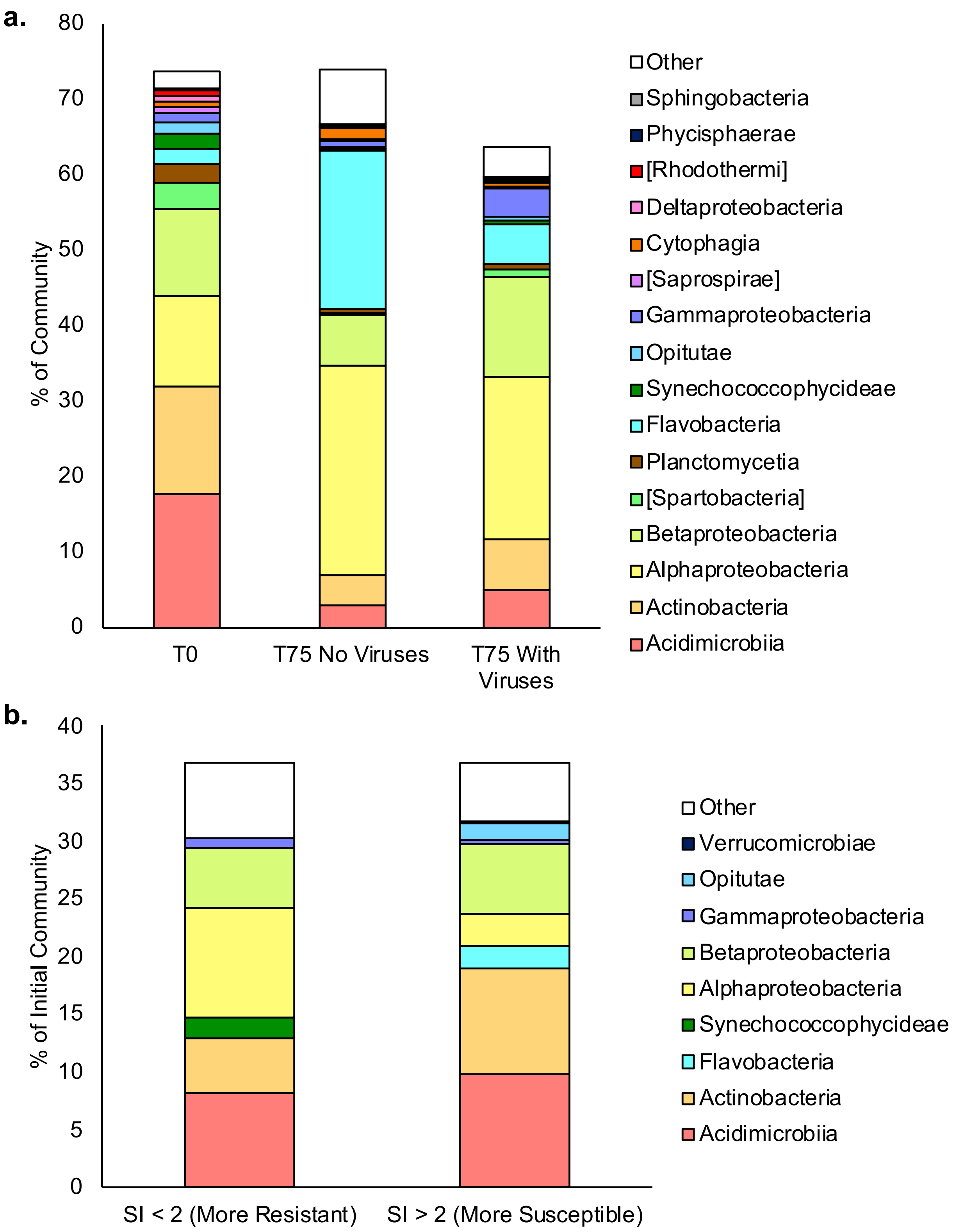


**Figure S8**. The impact of viruses on Chesapeake Bay summer bacterial populations. Bacterial communities sampled in July 2019 were incubated for 75 hours with (n = 6) or without (n = 6)

viruses. A) Bacterial community composition prior to and after incubation. Only those populations with observed growth (absolute abundance) during the incubation were included in the analysis. These populations comprised 74% of the initial community, 74% of the community after incubation without viruses, and 64% of the community after incubation with viruses. B) Community composition of putative resistant and susceptible bacterial populations in initial sample prior to incubation based on their growth in the presence or absence of viruses. Susceptibility index (SI) was calculated as (Population Abundance Fold Change Without viruses)/(Population Abundance Fold Change With Viruses). Higher values indicate greater susceptibility to viral-mediated mortality.

### SUPPLEMENTAL REFERENCES

1 Lane, D. in *Nucleic acid techniques in bacterial systematics* 115-175 (John Wiley and Sons: Chichester, UK, 1991).

2 Preheim, S. P., Perrotta, A. R., Martin-Platero, A. M., Gupta, A. & Alm, E. J. Distribution-based clustering: using ecology to refine the operational taxonomic unit. *Appl. Environ. Microbiol.* **79**, 6593-6603 (2013).

3 Klindworth, A. *et al.* Evaluation of general 16S ribosomal RNA gene PCR primers for classical and next-generation sequencing-based diversity studies. *Nucleic acids research* **41**, e1-e1 (2013).

4 Spencer, S. J. *et al.* Massively parallel sequencing of single cells by epicPCR links functional genes with phylogenetic markers. *The ISME journal* **10**, 427 (2016).

5 Preheim, S. P. *et al.* Surveys, simulation and single-cell assays relate function and phylogeny in a lake ecosystem. *Nature microbiology* **1**, 16130 (2016).
